# Ecologically pre-trained RNNs explain suboptimal animal decisions

**DOI:** 10.1101/2021.05.15.444287

**Authors:** Manuel Molano-Mazon, Yuxiu Shao, Daniel Duque, Guangyu Robert Yang, Srdjan Ostojic, Jaime de la Rocha

## Abstract

The strategies found by animals facing a new task are determined both by individual experience and by structural priors evolved to leverage the statistics of natural environments. Rats can quickly learn to capitalize on the trial sequence correlations of two-alternative forced choice (2AFC) tasks after correct trials, but consistently deviate from optimal behavior after error trials, when they waive the accumulated evidence. To understand this outcome-dependent gating, we first show that Recurrent Neural Networks (RNNs) trained in the same 2AFC task outperform rats as they can readily learn to use across-trial information both after correct and error trials. We hypothesize that, while RNNs can optimize their behavior in the 2AFC task without any *a priori* restrictions, rats’ strategy is constrained by a structural prior adapted to a natural environment in which rewarded and non-rewarded actions provide largely asymmetric information. When pre-training RNNs in a more ecological task with more than two possible choices, networks develop a strategy by which they gate off the across-trial evidence after errors, mimicking rats’ behavior. Population analyses show that the pre-trained networks form an accurate representation of the sequence statistics independently of the outcome in the previous trial. After error trials, gating is implemented by a change in the network dynamics which temporarily decouples the categorization of the stimulus from the across-trial accumulated evidence. Our results suggest that the suboptimal behavior observed in rats reflects the influence of a structural prior that reacts to errors by isolating the network decision dynamics from the context, ultimately constraining the performance in a 2AFC laboratory task.

## Introduction

In order to make good decisions in real life scenarios, animals are equipped with extensive implicit knowledge about the world. For instance, in foraging tasks, animals balance exploitation of the best alternatives with exploration of the less promising ones (Addicott et al. 2017), a behavioral pattern that seems imprinted in the brain (Daw et al. 2006; Blanchard and Gershman 2018; Chakroun et al. 2020) and that is present even when the difference in the alternatives values is large and stable (Vulkan 2000). These innate behaviors, that we call structural priors, shape the entire landscape of behavioral solutions available to the animal. However it is still unknown how to identify them and to which extent they influence the behavioral strategy adopted by animals in a given task.

Structural priors are thought to impact the behavior of animals in tasks organized sequentially into trials. For instance, animals’ responses are influenced by the history of sensory stimuli (Akaishi et al. 2014; Fischer and Whitney 2014; Akrami et al. 2018) and of previous responses and outcomes (Corrado et al. 2005; Lau and Glimcher 2005; Busse et al. 2011; Donahue, Seo, and Lee 2013; Abrahamyan et al. 2016; Urai et al. 2019). These sequential effects are suboptimal in most laboratory tasks in which trials are presented independently. However, they are highly prevalent and robust, which may indicate the existence of a hardwired circuitry that prevents animals from learning and implementing a more optimal strategy.

Many sequential effects are directly related to the outcome of the decisions made by the animal. Feedback has been shown to impact the bias towards the different options (Donahue, Seo, and Lee 2013; Abrahamyan et al. 2016; Urai et al. 2019), the speed of the subsequent responses (Rabbitt and Rodgers 1977) and the strategy used by subjects (Fusi et al. 2007; McDougle et al. 2016; Purcell and Kiani 2016b; Sarafyazd and Jazayeri 2019). It has been recently shown that rats are able to use the recent history of transitions after correct trials but consistently ignore it after error trials, following a suboptimal reset strategy (Hermoso-Mendizabal et al. 2020). These studies demonstrate that animals do not process success and failure merely as mirroring outcomes and that the neural mechanisms they trigger are qualitatively different (Lyon and Kuchling 2021). However, the structural prior underlying this asymmetry is still a matter of much debate (Baumeister et al. 2001; Alves, Koch, and Unkelbach 2017).

Recurrent Neural Networks (RNNs) constitute a useful tool to study the neural circuit mechanisms implementing the computations required to solve sequential tasks (Sussillo 2014; Barak 2017; Yang and Wang 2021). RNNs are typically allowed to adjust their connections during training to adopt the best strategy for the particular task, without any constraint imposed by preexisting knowledge (i.e. structural priors). However, this approach can potentially produce networks that use fundamentally different solutions from the ones used by animals, which are influenced by a plethora of structural priors. To overcome this mismatch between animals and RNNs, it has been recently suggested that networks should be pre-trained in more naturalistic tasks that induce the necessary priors in the networks before they face the laboratory task (Ma and Peters 2020; Yang and Molano-Mazón 2021). However, to our knowledge, no study has shown a successful example of the need of pre-training RNNs in order to replicate a sup-optimal behavior observed experimentally.

Here, we use RNNs to show that suboptimal history biases can emerge from brain circuits having evolved in environments with more naturalistic statistics than those characterizing laboratory tasks. We first conduct rat behavioral experiments to show that the suboptimal reset strategy adopted in a 2AFC task with serial correlations (Hermoso-Mendizabal et al. 2020) is a prevalent behavior across individuals and task variants. RNNs trained directly on the 2AFC task failed to replicate this behavior, outperforming the rats by fully exploiting the binary structure of the task. On the other hand, RNNs pre-trained in more naturalistic environments containing more than two alternatives replicate the reset strategy used by rats. Furthermore, as rats do, pre-trained networks maintain the trial history information after making a mistake, and use it again as soon as they made a correct choice. Finally, performing population analyses on the activity of the pre-trained RNNs, we identify the mechanism that allows them to cancel out the impact of trial-history on choice only after error trials. Our results suggest that the suboptimal strategy exhibited by rats in the 2AFC task could be the result of a hardwired structural prior that both guides and constrains their learning towards solutions consistent with more natural environments. Furthermore, our neural analyses provide specific, testable predictions regarding the neural mechanisms underlying the integration of information at different timescales. Finally, our work demonstrates that comparing animals’ behavior with that exhibited by RNNs may benefit from pre-training the networks in more ecologically relevant environments before testing them on the task of interest.

## Results

### Rats develop a robust suboptimal behavior in a 2AFC task with serial correlations

To investigate the extent to which rats utilize the information present in the statistical structure of the trial-to-trial sequence, we trained them in an auditory two-alternative forced choice (2AFC) task that included serial correlations (Hermoso-Mendizabal et al. 2020). In particular, the task was structured into trial blocks in which the probability that the previous stimulus category was repeated in the current trial, Prep, varied between high and low values (Prep=0.8 in repeating blocks and Prep=0.2 in alternating blocks; block size 80-200 trials; Fig. 1a and Supp. Fig. 13). Within each category, we parametrically varied the amount of evidence provided by each stimulus on a trial-by-trial basis, thus being able to build psychometric curves (Fig. 1b-c). Rats learned both to categorize the current stimulus and to infer Prep using the history of previous repetitions and alternations, what we called the transition history: within the repeating block, rats showed a tendency towards repeating their previous choice, whereas in the alternating block they tended to alternate (Fig. 1b). However, rats only displayed this transition bias after correct trials. After error trials, they consistently ignored such information and followed a strategy only based on the current stimulus (Hermoso-Mendizabal et al. 2020) (Fig. 1c). Importantly, this strategy, which we called reset strategy, is suboptimal in an environment presenting only two alternatives and in which contexts are stable (i.e. repeating/alternating blocks are long). A more optimal agent would, after an error trial, perform the simple counterfactual inference: “had I chosen the opposite side, I would have received a reward”; and use such inference to anchor the transition bias from the non-chosen side. For example, if the current estimate of the transition probability was to alternate, after an incorrect Right choice the agent should infer that the reward was present in the Left port and generate an alternation from there (i.e. Left→ Right), resulting in a rightward bias in the next choice (Fig. 1d). Because such a strategy implies a momentary reversal of the transition bias after an error trial, we call it the reverse strategy (Fig. 1e, gray area). Note that because the probability of an incongruent transition is relatively high (e.g. repetitions in the alternating block occur with probability Prep=0.2), a single error should not be interpreted as a block change which occurs with a much lower probability (P<1/80).

**Figure 1.**
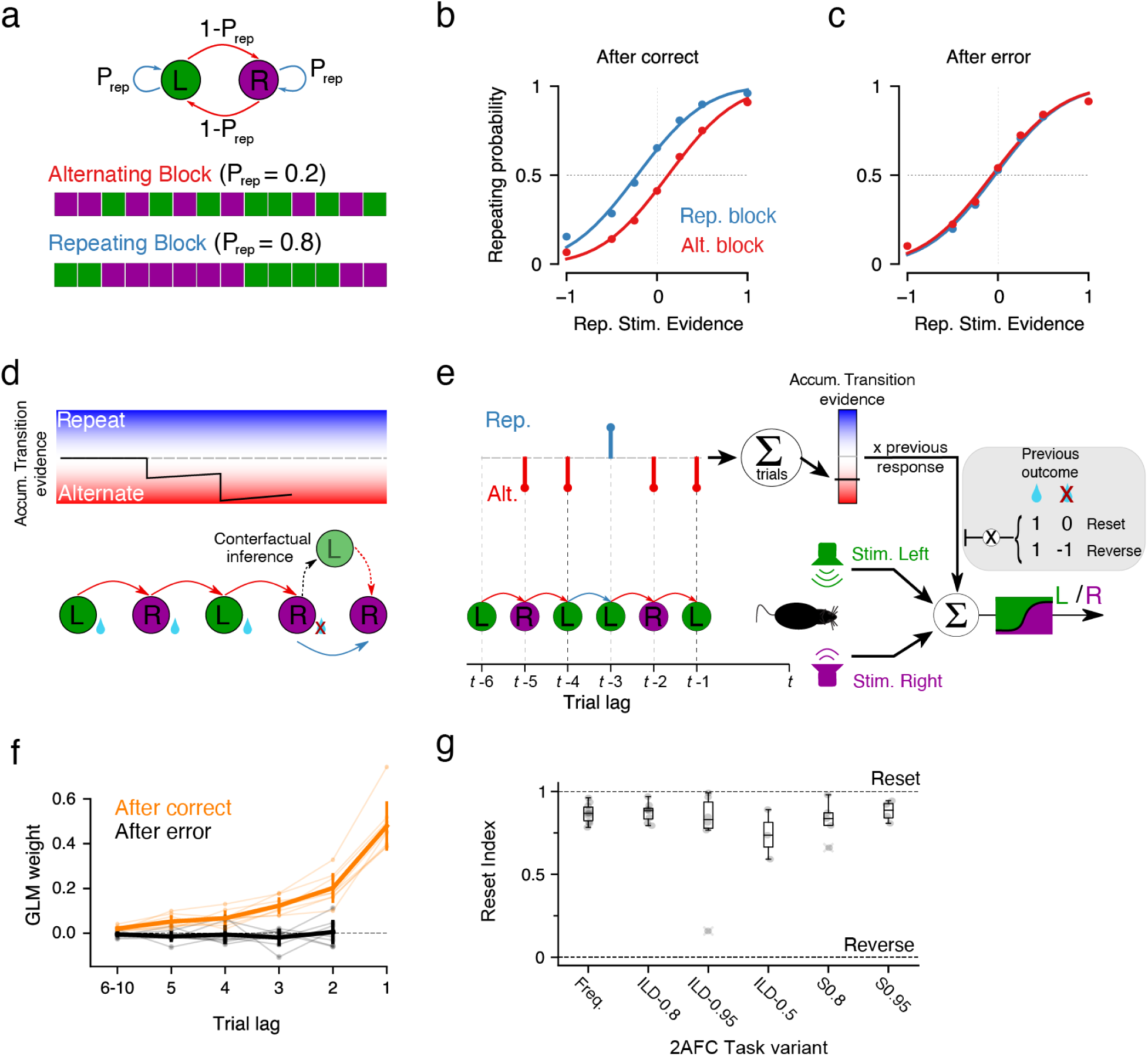
Rats develop a history bias only after correct trials. **a)** The trial sequence of stimulus categories in the 2AFC task is generated by a two-state Markov chain parametrized by the repeating probability P_rep_ (top) which is varied in Repeating (P_rep_=0.8) and Alternating trial blocks (P_rep_=0.2) (bottom). Block length was set between 80 and 200 trials. **b-c)** Average psychometric curves showing the probability of repeating the previous choice as a function of the stimulus evidence supporting the repetition (rat Group ILD-0.8, n=8 rats). Curves were computed separately for trials after correct **(b)** and after-error **(c)** choices, and for the repeating (blue) and alternating (red) blocks. Dots represent the data and lines are logistic fits. The shift of the curves shows the repeating (blue) or alternating bias (red) caused by trial history. **d)** Example sequence of choices made by an optimal agent (bottom), together with the trace of the accumulated transition evidence (top). After an error the agent infers the rewarded side (black arrow) and uses the accumulated alternating evidence to bias its next response towards repeating the previous choice (effectively reversing the alternating bias). **e)** Rats’ responses are modeled using a Generalized Linear Model that combines the current stimulus evidence (bottom) and the history of transitions, i.e. previous repetitions (blue) and alternations (red). The weighted sum of transition history provides the accumulated transition evidence (color bar) which captures the current tendency to repeat or alternate the previous rewarded side. Shaded area shows the effect that the previous outcome has on the contribution of the transition evidence, both in the reset strategy exhibited by rats and in the reverse strategy expected from an ideal observer. **f)** GLM weights of previous correct transitions T^++^ (i.e. formed by two correct choices) computed after correct (orange) and after error (black) trials; mean and std. dev. from rat Group ILD-0.8 (n=8, thick traces) and individual animals (light traces). **g)** Reset Index (see methods) for different rat groups performing different task variants. Group Freq.: frequency discrimination task, n=10 (Hermoso-Mendizabal et al. 2020); Group ILD-0.8: intensity level discrimination task with P_REP_ = 0.8, n=8; Group ILD-0.95: ILD task with P_REP_ = 0.95, n=6; Group ILD-Uncorr: ILD task with uncorrelated sequences, (i.e. P_REP_ = 0.5), n=3; Group S0.8: silent task (without any stimuli) and P_REP_ = 0.8, n=5; Group S0.95: silent task with P_REP_ = 0.95, n=4. The reset strategy implies RI∼1 whereas in the reverse strategy RI∼0.

To quantify the extent to which the behavior of individual rats followed the reset or the reversal strategies, we modeled the rats’ choices using a Generalized Linear Model (GLM) (Lau and Glimcher 2005; Abrahamyan et al. 2016; Braun, Urai, and Donner 2018). The GLM assumes that the rats’ choices are the result of a linear combination of various task variables such as the current stimulus evidence or the previous choices and transitions together with their outcomes (Fig. 1e and Supp. Fig. 1) (see Methods). This analysis allows us to quantify the contribution of the past correct transitions T^++^ (those formed by two consecutive correct choices) to the rat’s current choice. To uncover the potentially different strategies followed by rats depending on the outcome in the previous trial, we separately fitted their choices after correct and error trials with two different models. Thus, the model fitting after-error choices should yield vanishing weights associated to the previous transitions for the reset strategy, while showing negative weights for the reverse strategy (Fig. 1e, inset). We found that rats consistently utilized the transition history after a correct trial and ignored it after an error (Fig. 1f). We used the fitted T^++^ transition weights from after correct and after error trials to define a Reset Index (RI), which was zero for perfectly symmetric kernels (i.e. reverse) and approached one as the after-error kernel vanished (i.e. reset; see Methods). The RI was close to one (mean±SD 0.83 ±0.06) for almost every animal tested in different variants of the 2AFC task (Fig. 1g and Supp. Fig. 2), in which we changed the stimulus feature to be categorized (frequency or intensity) and the repeating probability (Prep = 0.8-0.2, 0.95-0.05 and 0.5-0.5, i.e. no serial correlations). Furthermore, we trained rats in a variant of the task with no stimuli (Duque and de la Rocha 2022) (Fig. 1g, Group S0.8, n=5), and with extremely predictable repeating and alternating sequences (Prep = 0.95-0.05, Group S0.95, n=4) with the aim of making the task conceptually simpler and thus the pattern of transitions more evident. Despite this major simplification, rats still displayed a robust reset strategy. Moreover, rats also discarded any information provided by transitions containing incorrect choices (i.e. *T*^+−^, *T*^−*+*^ and *T*^−−^ transitions) (Supp. Fig. 3) (Hermoso-Mendizabal et al. 2020). Importantly, as with the reset strategy, this disregard of any information involving error trials is not optimal, since in the 2AFC task all types of transitions are equally informative (Supp. Fig. 4). This robust behavior further supports the hypothesis that there exists a fundamental difference between the strategies followed by rats when dealing with correct and error outcomes.

### RNNs learn to fully leverage trial-history information

What causes the reset strategy observed in rats? One possibility is that the counterfactual inference needed after error trials constitutes a complex computation that is difficult to implement in a neural circuit. To test this hypothesis, we trained Recurrent Neural Networks (RNNs) in the 2AFC task. RNNs were presented with noisy stimuli that provided evidence for each of the choices and that were organized into Repeating and Alternating blocks, as in the rats experiments (Fig. 2a). In order to test the ability of RNNs to perform counterfactual inference, we trained them using Reinforcement Learning (RL) (Z. Wang et al. 2016; Sutton and Barto 2018), by which networks only receive feedback in the form of a scalar (the reward), without being explicitly told the correct answer at each timestep, as it is done in Supervised Learning techniques (Werbos 1990). Further, we provided the networks with the reward and action from the previous timestep, which allows them to be more adaptive to the different contexts of a task (J. X. Wang et al. 2018). At each timestep the output of the networks could be to fixate, to choose left or to choose right (Fig. 2a; see Methods).

**Figure 2.**
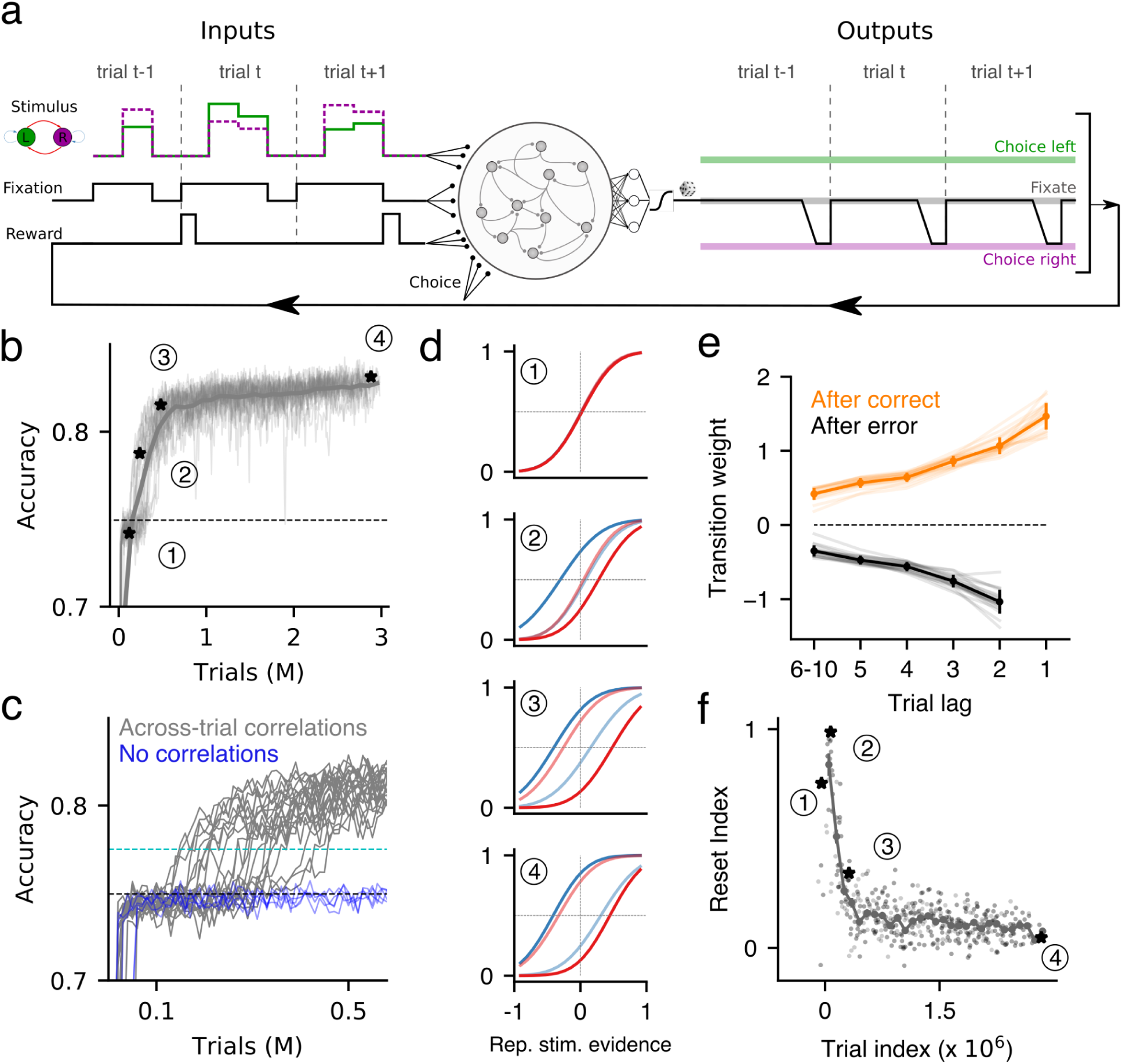
Behavior of RNNs trained directly on the 2AFC task. **a)** LSTM networks containing 1024 units were trained using Reinforcement Learning (Z. Wang et al. 2016). At each timestep, the networks received as input a fixation cue, two inputs corresponding to the two stimuli (left/right), and the reward and action at the previous timestep; in turn, at each timestep networks had to fixate, go left or go right (see Methods). The example shows a hypothetical sequence in which the network (middle) experiences 3 alternating trials (left) and chooses the right side at the end of each of them (right). **b)** Accuracy across training of 16 RNNs (thin traces). The thick, dark line corresponds to the median accuracy. Dashed line corresponds to the accuracy of an agent that only uses the current stimulus evidence and not the transition history to make a decision (see Methods). Black asterisks and numbers correspond to different periods of training for which we plotted the psychometric curves in panel d. **c)** Expanded view of panel b showing the period during which RNNs learn to use the transition history. Aha-moments were detected by setting a threshold on the accuracy (dashed cyan line). Blue traces show the accuracy of RNNs trained on a 2AFC task with no across-trial correlations (i.e. P_REP_ = 0.5). **d)** Psychometric curves as the ones shown in Fig. 1b-c, for the different training periods indicated in panel b. **e)** Transition kernels as the ones shown in Fig. 1f corresponding to trained networks (i.e. point 4 in panels a, d and f). Error-bars correspond to standard deviation. **f)** Reset Index values for the same networks shown in panel b, when tested in the 2AFC task at different stages of training (light dots). Values are aligned to each network’s aha-moment (see c). Dark dots correspond to the median reset index across individual networks. Error-bars correspond to standard error.

RNNs quickly learned to integrate the stimuli to inform their decision, reaching an accuracy comparable to that of an agent that integrates the stimulus perfectly but acts independently of previous history (Fig. 2b, dashed line). After following this perfect-integrator strategy for a variable period of time, networks underwent an aha-moment and started exploiting the information provided by the transition history, hence further improving their accuracy (Fig. 2b, c). This shift in strategy occurred relatively fast and in a few thousand trials the accuracy of the networks reached a relative plateau in which all RNNs used both the stimulus evidence and the transition history (Fig. 2b-d). Importantly, at the beginning of this plateau, the behavior was close to the reverse strategy, with the transition bias being present both after correct and after error trials (Fig. 2b-d, point 3). Along this final phase of training, the reverse strategy was refined with the transition bias after correct and error trials becoming more pronounced and symmetric (Fig. 2b-d, point 4). Thus, RNNs trained directly on the 2AFC task learned to leverage both the stimulus evidence and the transition history information. We performed the same GLM analysis carried out in rats (Fig. 1e-f) and found that, after training, RNNs showed a strong contribution of the past correct transitions T^++^ in both after correct and after error choices (Fig. 2e). This implies the networks reached a very low Reset Index along the training (Fig. 2f). Importantly, RNNs trained in the same task but with zero across-trial correlations did not undergo any aha-moment but maintained the performance of the agent only using stimulus evidence (Fig. 2c blue). Consistently, they showed no contribution of previous transitions on choice (Supp. Fig. 5). Thus, when possible, RNNs capitalized on the predictability of the sequence and showed a transition bias both after correct and after error responses. That simple RNNs could fully exploit the symmetry of the 2AFC task and adopt a reversal strategy (Fig. 1d) suggests that rats’ reset strategy was not caused by a limit of their computational capacity but by some more fundamental factors we aim to explore next.

### The reset strategy is adaptive in environments with more than two alternatives

If the reverse strategy can be implemented in a small neural circuit, why do rats reset after making a mistake? We reasoned that a fundamental difference between the training of our rats and our RNNs is that, while the former face the learning of the task with an extensive background knowledge about the statistical structure of the world, RNNs start as a blank slate whose wiring can be fully optimized to perform the 2AFC task. Therefore, the reset strategy could have originated from evolution as an adaptation to more naturalistic environments, giving rise to a structural prior that hinders the rats from fully exploiting the structure of the 2AFC task. We hypothesize that one key difference between the 2AFC task and naturalistic environments is the number of available alternatives. As explained above (Fig. 1d), in any 2AFC task subjects have always implicit access to the identity of the correct alternative, independently of the outcome of their decision. Thus, correct and incorrect answers provide the same exact amount of information about the environment. This is no longer true when we increase the number of alternatives to more than two: in a task with N>2 alternatives, information about the reward location is ambiguous after making an incorrect choice, because it could be any of the other N-1 alternatives. This means that exploiting serial correlations after error trials, although optimal for any N, only provides a 1/N decreasing benefit (i.e. by discarding one choice out of N). Therefore one would expect that, for sufficiently large Ns, RNNs will adopt the near-optimal reset strategy that only exploits serial correlation after rewarded responses (Fig. 3b). Can this reset strategy, presumably adopted for large N environments, form a structural prior that constrains learning at any N or are networks able to adapt their strategy depending on N?

**Figure 3.**
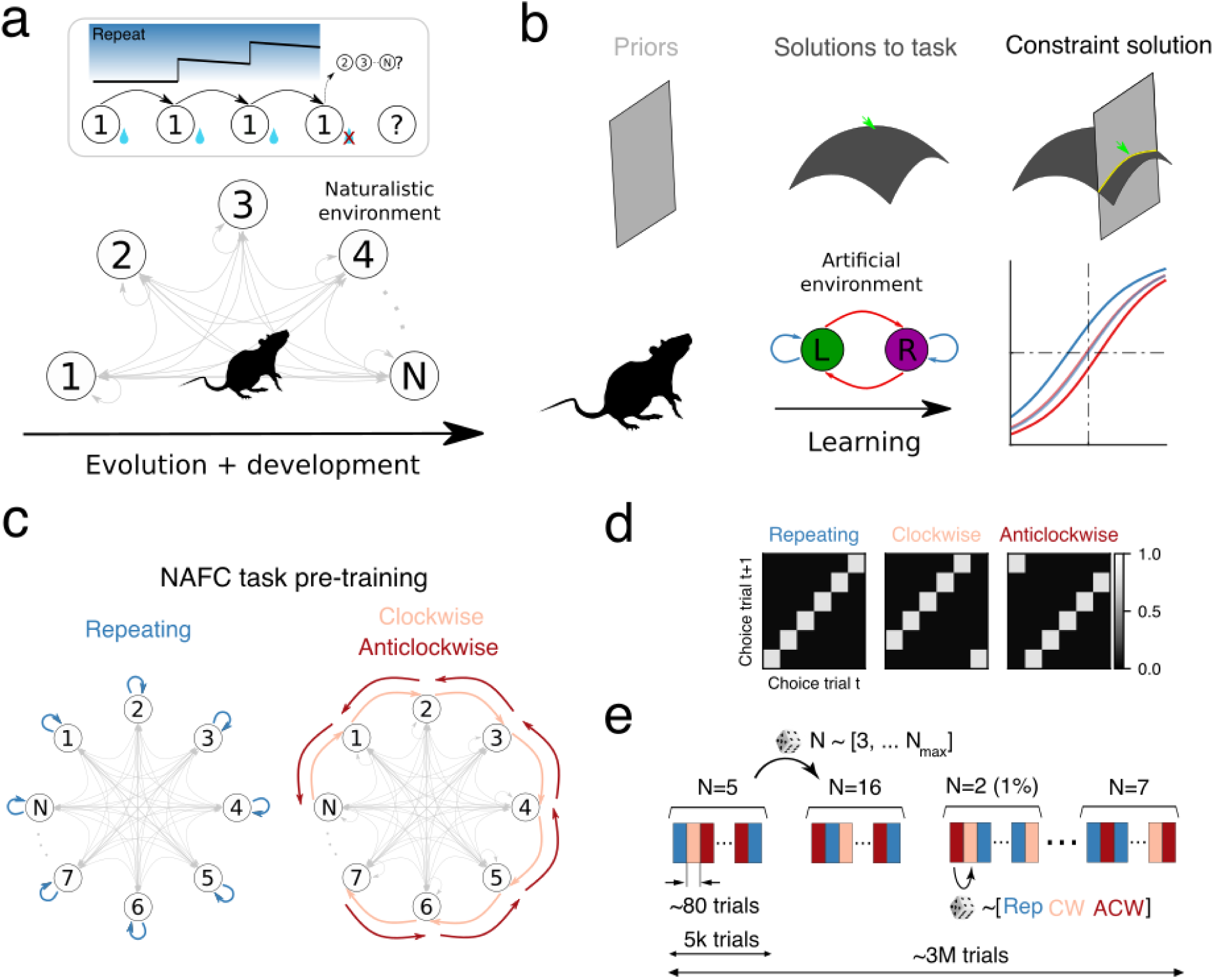
The reset strategy naturally emerges in an environment presenting more than two alternatives. **a)** Rats have evolved in naturalistic environments with more than two alternatives (bottom) in which a single error prevents subjects from predicting the next state of a sequence despite knowing the transition probabilities (top). **b)** Cartoon describing how the priors imposed by evolution and development in naturalistic environments (left column) affect the animal’s capacity to find the optimal solution in the 2AFC task (middle columns, green arrow); the final solution (right column) constitutes a compromise between the priors and the task’s optimal solution. **c-d)** In the N-AFC task the across-trial stimulus category sequence is organized into contexts **(c)** with distinct serial correlations, each parametrized by a transition matrix **(d)**. We pre-trained the networks using repeating, clockwise and anticlockwise contexts, each one having a most likely sequence (colored arrows) and less frequent transitions (gray arrows). **e)** During pretraining in the NAFC environment, block length was random (avg. duration of 80 trials) and the number of choices N was changed every 5k trials. The percentage of trials with N = 2, i.e. the 2AFC task, was set to 1%.

### Pre-training RNNs in a more ecological task recovers the reset strategy

To answer this question we embedded the 2AFC task in a more general NAFC environment presenting a variable number of alternatives N that was in general larger than two (Fig. 3c, Supp. Fig. 15a-d and Methods). With this simplified setup, we seeked to emulate the existing tension between the structural prior, modeled as the general strategy used when N can be large, and the rat’s necessity to learn a new, artificial laboratory task represented by the N=2 case. Thus, RNNs trained in these conditions, called pre-trained RNNs, were pushed to find the best solution for the 2AFC task within the realm of solutions set by the NAFC environment. By generating a sequence of rewarded states using non-homogeneous transition probability matrices, the NAFC environement modeled the spatiotemporal correlations commonly found in natural environments. In particular, trials were organized into a repeating, a clockwise and an anticlockwise context, each defined by a different transition matrix and presented randomly (Fig. 3c-e). The maximum number of alternatives was fixed to N_max_ and the number of available alternatives, N, varied randomly between 2 and N_max_ every 5k trials (Fig. 3e). Independently of the value of N_max_, blocks with N=2 constituted a fixed, small fraction (1%) (Fig. 3e).

To characterize how RNNs learned to exploit serial correlations, we computed the accuracy in the NAFC of pre-trained networks on trials with zero stimulus evidence. From early stages of the training, the average accuracy was above chance for all values of N (Fig. 4a), demonstrating that networks quickly developed and used a transition bias regardless of the number of alternatives. However, in after-error trials, RNNs exhibited chance accuracy throughout the pre-training suggesting that the NAFC environment with multiple Ns precluded networks from developing the reverse strategy at any value of N (Fig. 4b).

**Figure 4.**
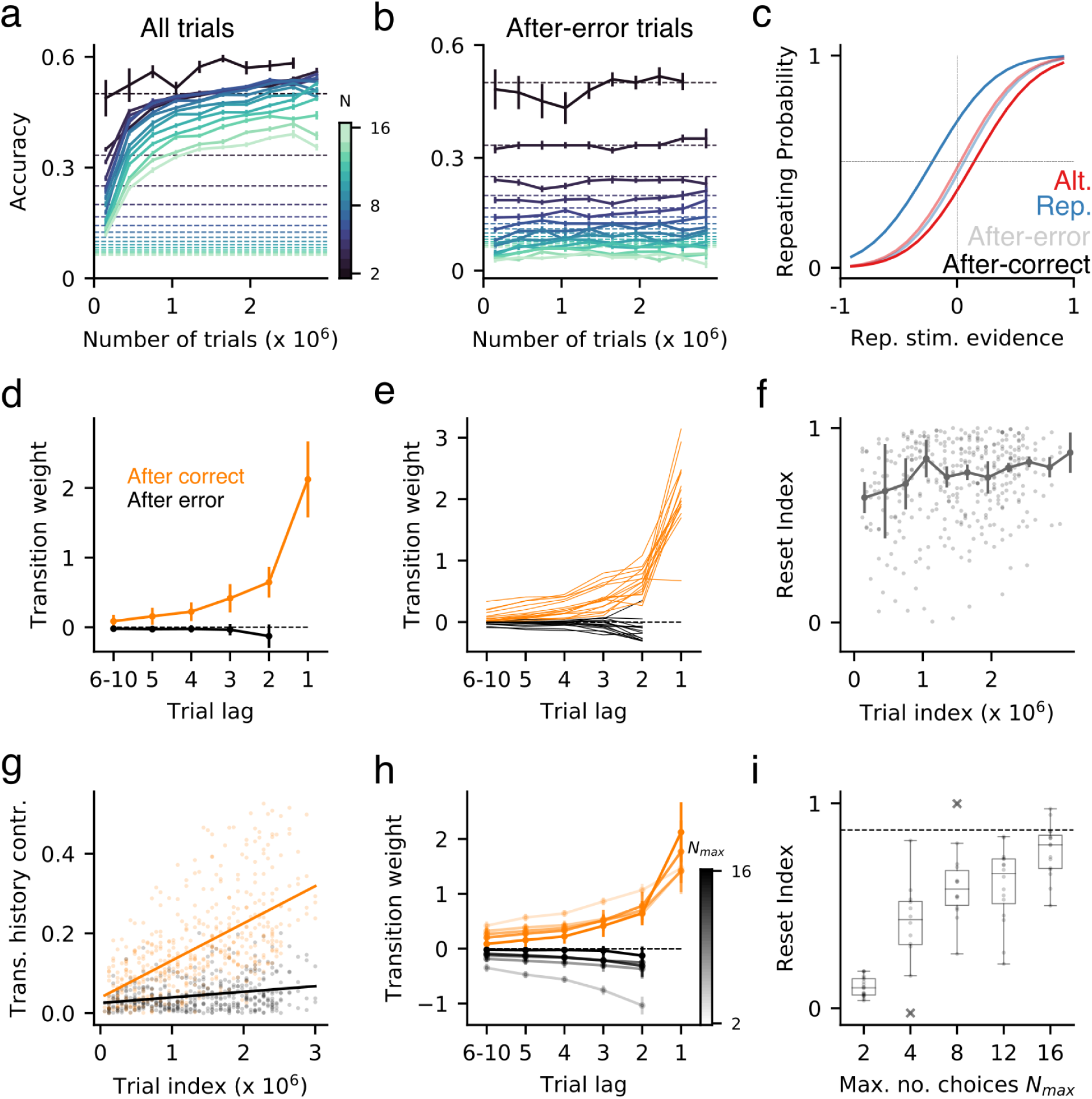
Behavior of networks pretrained in a 16-AFC task. **a)** Median accuracy (n=16) for trials with zero stimulus evidence conditioned on the number of available alternatives N=2, 3, 4, 5, 6, … , 16 (see colorbar). Dashed lines indicate the chance level (1/N) corresponding to each value of N. **b)** Same as in panel a for trials following an incorrect choice that is congruent with the transition probabilities of the current context. Color-code as in panel a. **c)** Psychometric curves obtained for each block (see color code) when testing a pre-trained RNN on the 2AFC task, computed in after-correct (dark) and after-error (light) trials. **d)** Average after-correct and after-error transition kernels for pre-trained RNNs tested in the original 2AFC task. **e)** Same as d for individual networks. **f-g)** Median Reset Index **(f)** and contribution of transition history **(g)** in the 2AFC task across pre-training. Points show individual RNNs. Transition history contribution is computed after correct (orange) and after error (black) trials as the absolute value of the sum of the individual weights at different lags (see Methods). Lines correspond to linear fits. **h)** Average transition kernels obtained from networks pre-trained with different maximum number of alternatives N_max_ (see colorbar). **i)** Reset Index versus N_max_. Dots correspond to individual networks. Dashed line indicates the mean Reset Index obtained across all rats and task variants. Error-bars correspond to standard error in panels a, b and f and standard deviation in panels d and h.

We next quantified in more detail the extent to which the structural prior induced by the NAFC environment conditioned the strategy found by the RNNs in the 2AFC task. We characterized the learned behavior by freezing the connectivity and testing the RNNs in the 2AFC task, thus preserving the wiring sculptured by the pre-training. The pre-trained networks followed the reset strategy observed in the rats. This can be seen in the difference between the psychometric curves obtained after correct and after error trials (Fig. 4c) and was confirmed by the vanishing transition weights obtained from the GLM fit after error trials (Fig. 4d, e). The reset strategy was steadily developed through pre-training suggesting that it was not a partial solution found due to incomplete training (Fig. 4f-g). The asymmetry between after-correct and after-error transition weights increased as the maximum number of alternatives Nmax increased (Fig. 4h), which made the Reset Index rise accordingly reaching values similar to those found in rats for Nmax=16 (Fig. 4i). Furthermore, pre-trained networks not only ignored transition history after error trials, but for large Nmax they mostly disregarded transitions containing at least one error, just as rats do (compare Supp. Fig. 1 and Supp. Fig. 6). Therefore, although the more optimal behavior was to develop a reversed transition bias after errors, particularly for low Ns, RNNs pre-trained in the NAFC environment adopted a general reset strategy that was near optimal but common for all Ns.

The reset behavior stemmed from two features of the pre-training: the information asymmetry between after-correct and after-error trials in the NAFC environment and the appropriate balance between the learning of the 2AFC and the contextual influence of the NAFC. To show the key role of these factors, we first pre-trained RNNs in a variant of the NAFC environment in which networks received the identity of the correct alternative, independently of the outcome of their response. Having access to this information, networks learned to make use of the transition history both after correct and error trials, showing a clear reverse strategy (Supp. Fig. 7a). Second, we checked that the reset strategy adopted in the 2AFC was not caused by the RNNs not having experienced a sufficient number of N=2 trials during the pre-training. When we varied the proportion of N=2 trials, pre-trained RNNs showed a robust reset behavior for a substantial range of values 0-2.5% and only when the proportion reached 25% pre-trained RNNs systematically adopted the reversal strategy (Supp. Fig. 7b). On the other hand, although pre-training prevented the adoption of the reverse strategy in the 2AFC task, it also facilitated a faster learning to exploit, at least partially, the across-trial serial correlations: pre-trained RNNs needed many fewer 2AFC trials to reach asymptotic accuracy than networks trained directly in the 2AFC (Supp. Fig. 7c). By contrast, removing the serial correlations from the NAFC (for N>2), promoted a memoryless trial-history prior that burdened pre-trained RNNs from learning transition biases (Supp. Fig. 7c). Thus the pre-training in the NAFC environment both constrained and helped the learning of the 2AFC task, guiding it towards a more consensual solution.

### Pre-trained RNNs maintain the transition history information after an error

Having shown that pre-trained networks display the reset strategy after error choices, we next aimed to understand the dynamics of the transition bias beyond this reset. Do pre-trained networks recover part of the accumulated transition evidence after an error trial or does the inference of the transition probabilities start up *de novo*? The adaptive behavior in the NAFC, as in any environment with more than two alternatives and relatively stable contexts, is to transiently gate off the use of the accumulated transition evidence after an error but recover its use after the next correct choice. For example, if the network is in a repeating block and makes an error, then it should go unbiased until making a correct choice, after which it should resume repeating because, most likely, it remains in a repeating context. To test whether pre-trained RNNs exhibited such a gate-and-recovery behavior, we investigated the across-trial dynamics of the latent transition bias matrix T*_ij_*(t) (Fig. 5b). This matrix quantifies the average probability that, having selected choice *i* in trial *t*-1, the networks select choice *j* in trial *t* in response to a stimulus with zero evidence. As an example, we computed T*_ij_*(t) throughout a sequence composed of three correct responses forming a pattern congruent with the anticlockwise context (ACW), followed by an error and then a correct response (Fig. 5c). After the first correct ACW transition, the networks showed a clear tendency to reproduce the ACW transition in the next trial, a tendency that got more pronounced after the second correct ACW transition (Fig. 5b, left panels). An error response afterwards completely flattened the transition matrix, reflecting the reset strategy. However, a single correct response after the error, recovered the networks’ ACW tendency, evidencing that the transition history accumulated previous to the error had not been erased (Fig. 5b, inset in the rightmost panel). Interestingly, the transition probabilities at recovery also show a weaker tendency towards the other two possible contexts (CW and REP, in Fig. 5), showing that the decision of the networks was influenced by both the recent transitions and by knowledge of the existing contexts learned during the pre-training. The networks were therefore able to update and maintain an internal representation of the accumulated transition evidence, transiently gating it off after errors and recovering it after the next correct choice (Fig. 5a-b, Supp. Fig. 8a-b).

**Figure 5.**
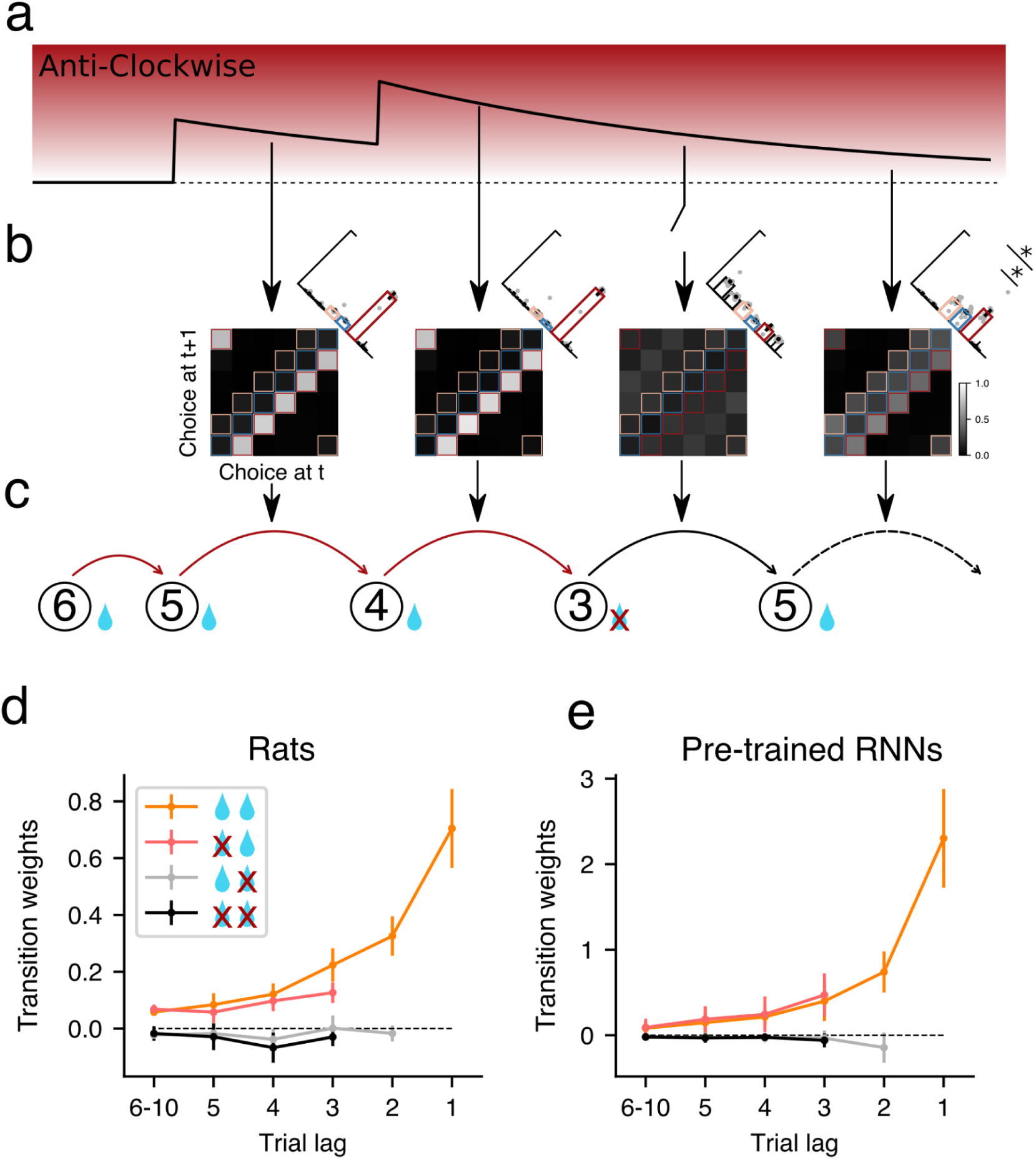
Pre-trained networks show the gate-and-recovery dynamics observed in rats after an error followed by a correct response. **a-c)** Schematic illustrating the across-trial dynamics of a latent variable encoding the transition probabilities throughout a sequence containing three correct responses following an ACW pattern, followed by an error and then a correct response (panel c shows an example of such sequence). Notice that the occurrence of an error does not reset this latent probability (a) but it gates off its impact onto the next choice (see broken arrow). The evolution of the transition bias matrix throughout the sequence (b) was numerically computed using pre-trained RNNs (N_max_=16, RNNs are tested with N=6). Inset: histograms showing the probability of making a Repeating (blue), Clockwise (pink) and Anticlockwise (red) choice. In the last trial, the histogram shows a higher probability (p<0.002, paired t-test) to make an ACW than a Rep or CW choice. **d-e)** Average *T*++ transition kernel obtained in the 2AFC task when the GLM is fitted independently depending on the outcome of the last two trials for rats **(d;** n=10) and for the pre-trained RNNs **(e**; n=16**) (**see inset; e.g. a crossed water drop followed by a drop represents that *t-2* was incorrect and *t-1* was correct). Because the weights refer to T++ transitions, only when trials *t-1* and *t-2* are rewarded the kernel contains weights at all lags (i.e. -1, -2, … ; orange).

We then checked that the gating of the transition evidence was specific to after-error trials. First we confirmed that the transition bias was present after unexpected but correct responses that broke the context’s transition pattern (Supp. Fig. 8c-d). Second, we checked whether the stimulus evidence in the previous choice affected the transition bias in the current trial, as predicted by some normative models (Lak, Okun, et al. 2020; Lak, Hueske, et al. 2020). To do so, we selected the trials preceded by each of the sequences made of four correct choices (e.g. LRLL) and compared the transition bias for trials following an easy choice (i.e. strong stimulus evidence) versus those following a hard choice (i.e. weak stimulus evidence). We found that, for both rats and networks, the amount of stimulus evidence in the previous trial did not affect the transition bias, which only depended on the history of previous choices and outcomes (Supp. Fig. 9). Thus, the only modulation of the transition bias we could identify was the gating caused by the previous outcome.

The adaptive gate-and-recovery behavior developed by pre-trained networks in the NAFC percolated into the 2AFC task mimicking the behavior found in rats (Hermoso-Mendizabal et al., 2020). To show this, we computed the T++ transition kernel separately from trials conditioned on the outcome of the last two trials, giving rise to four conditions: after correct-correct, after error-error, after correct-error and after error-correct. After an error, the kernel weights were close to zero in both rats and pre-trained RNNs (Fig. 5d-e) consistent with previous analysis (Fig. 1f and 4d-e, respectively). However, after a correct response following an error, the weight of previously accumulated transitions (trial lags -3, -4, …) was recovered, indicating that those past transitions had again the same impact on choice as if there have not been any error (i.e. their weights were the same as after two correct responses). Thus, networks pre-trained in the NAFC reproduced the gate-and-recovery strategy displayed by rats in the 2AFC task, supporting the idea that their behavior is constrained by a structural prior deeply founded in the asymmetry between correct and error responses characteristic of natural environments.

### Neural mechanisms underlying the encoding of transition bias and the after-error reset

Next we studied the neural mechanisms underlying the transition bias and its reset after error trials. We reasoned that the underlying computation consisted of three stages: (i) integrating the transition history to estimate the current context; (ii) combining the estimate of context with information on the previous choice to infer the relevant transition bias; (iii) integrating the transition bias with the current stimulus to generate the choice. Our goal was to specifically determine which of these stages were impacted following an erroneous choice, leading to the after-error reset. We therefore performed linear decoding analyses of population activity during the testing on the 2AFC task, and contrasted after-correct and after-error trials in pre-trained RNNs with networks trained directly in the 2AFC task (2AFC-trained networks).

We first investigated to which extent networks encoded latent information about the current context in the population activity. For that we selected trials following several correct transitions in which inferring the context was easiest for the network (*clear context trials*; see Methods) and focused our analysis on the activity at fixation period right before the stimulus was presented. We trained a linear Support Vector Machine (SVM) to identify the dimension in the activity space along which the repeating and alternating contexts were best separated (Fig. 6a) (see Methods). When projecting the activity of all trials along this dimension, we found that the networks could correctly infer the current context for most trials (Fig. 6b). Importantly, this was also true for after error trials (mean ± std.dev. AUC=0.87 ± 0.04 for after correct trials; AUC=0.75 ± 0.06 for after error trials; n=16 networks), which supports the network’s gate-and-recovery behavior (Fig. 5e) by showing that the context encoding was not reset after errors. We found a qualitatively similar context encoding in 2AFC-trained networks (AUC=0.90 ± 0.05 for after correct trials, AUC=0.82 ± 0.05 for after-error trials) (Supp. Fig. 10a-b). We also observed a sharp update of context encoding at block change (Fig. 6c), which is consistent with the quick updating of the transition bias matrix (Supp. Fig. 8), and allow the networks to quickly adapt to block changes at the cost of being more sensitive to spurious fluctuations in the sequence of transitions (see occasional fluctuations in (Fig. 6b)).

**Figure 6.**
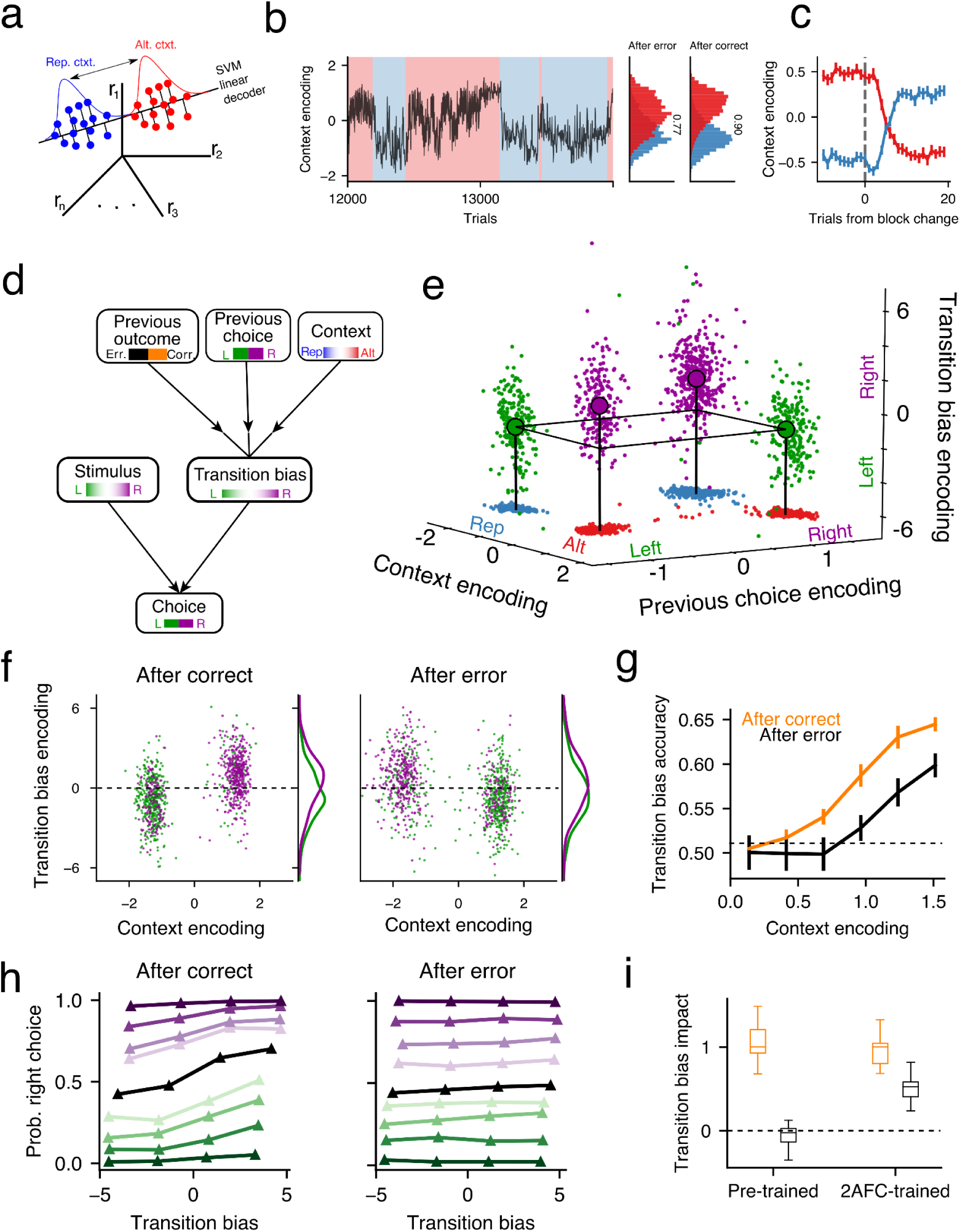
Network mechanisms underlying context and bias encoding in pre-trained networks. **a)** Schematic illustrating the population analysis. An SVM is trained on *clear context trials* that follow several correct repetitions or alternations to determine the direction in activity space that best separates the two contexts (black line). The projection onto this axis provides the trial by trial context encoding. **b)** Dynamics of context encoding at fixation time across 1000 trials spanning several blocks. **Right:** histograms for repeating (blue) and alternating (red) blocks obtained from after-error and after-correct trials, show that context encoding is highly accurate (AUC= 0.77 for AE and 0.90 for AC). **c)** Average context encoding aligned at repeating-to-alternating (red) and alternating-to-repeating block changes (blue). **d)** Schematic showing the relevant variables employed in the transition bias computation. The transition bias requires the integration of the estimated context, and the previous choice and outcome. The resulting bias is then combined with the stimulus to yield the final choice. **e)** Transition bias encoding as a function of the encoding of the context and the previous choice, shown for clear context trials. To visualize the accuracy of the bias prediction, dots are colored according to the most likely upcoming stimulus category (i.e. after an X correct choice, the new stimulus category is X in repeating blocks and Y in alternating blocks). Red and blue dots show the projection of each trial onto the (context, prev. choice) plane. Stems illustrate the center of mass of each of the four conditions. A level line with constant bias is shown to ease the comparison (black line). **f)** Transition bias encoding versus context encoding for after-correct (left) and after-error trials (right). Only clear context trials preceded by a Left choice are shown. Dots are colored according to the true upcoming stimulus category. Each cluster shows around 20% misclassified trials corresponding to *incongruent* transitions (e.g. a repetition in an alternating block). Transition bias histograms illustrate the accuracy of the transition bias predicting the upcoming stimulus (AUC=0.69 and 0.62 for after-correct and after-error trials). **g)** Transition bias accuracy versus context encoding computed separately for after-correct and after-error trials. Accuracy was calculated by categorizing the encoded transition bias and comparing this prediction with the true upcoming stimulus category. **h)** Psychometric curves showing the network’s choice as a function of the internal encoding of the transition bias for different values of the stimulus evidence in after-correct (left) and after-error (right) trials. Color code reflects stimulus evidence (darker colors show stronger evidence; black corresponds to zero evidence). **i)** Normalized slopes of the psychometric curves shown in (h) for zero stimulus evidence for pre-trained (left) and 2AFC-networks (right) for after-correct (orange) and after-error (black) trials. Slopes were normalized by the mean after-correct slope, separately for each type of RNN. Panels b-f,h show the same example pre-trained RNN whereas panels g, and i show the statistics from n=16 RNNs for each group. Dots in g shows the average across networks and error bars show s.e.m.

The internal representation of the context must be combined with the previous choice and previous outcome in order to compute the transition bias (Fig. 6d). Because this bias reflects the network’s prediction about the upcoming stimulus category, we trained a decoder to predict the following stimulus category from the network activity during the fixation period using clear context trials (see Methods). The readout of this decoder was used as a proxy of the transition bias. When representing the transition bias as a function of the context and the previous choice encoding, we found that, in clear context after-correct trials, the network gave rise to the appropriate bias in each of the four combinations of context and previous choice (e.g. previous left + alternating context = bias towards right) (Fig. 6e). We then assessed the dependence of the transition bias on the previous outcome. Unexpectedly, the pre-trained RNN slightly reversed the bias after errors, meaning that the upcoming stimulus category could in principle still be predicted, although with less accuracy than after correct choices (Fig. 6f-g, Supp. Fig. 11). The 2AFC networks, on the other hand, fully reversed the transition bias after errors and maintained an accurate prediction about the upcoming stimulus independently of previous outcome (Supp. Fig. 10c).

How can pre-trained RNNs show reset behavior after errors when they internally have a relatively accurate encoding of the upcoming stimulus (Fig. 6f-g)? We reasoned that the dependence of the final network’s choice on the magnitude of transition bias was probably different depending on the previous outcome. To test this, we computed psychometric curves as a function of the internal representation of the bias and the stimulus evidence. We found that the networks’ choice was strongly determined by the magnitude of this bias after correct trials but not after error trials, whereas the impact of the stimulus was comparable in the two conditions (Fig. 6h). This was also true when we separately trained the transition bias decoder using only after-error trials (Supp. Fig. 12). This implies that the previous outcome was used by the network as a gating signal to switch its decision dynamics from a bias-dependent categorization of the stimulus in after-correct trials to a categorization only based on the stimulus evidence in after-error trials. In contrast, in the networks trained directly on the 2AFC the transition bias encoding had comparable impact on choice in both types of trials (Fig. 6i and Supp. Fig. 10d). Therefore, pre-training in the NAFC environment endowed networks with a specific mechanism to decouple the stimulus categorization in after-error trials from the information accumulated in the transition bias, hence producing the gating strategy observed in their behavior.

## Discussion

We have investigated the causes for the suboptimal strategy displayed by rats in a 2AFC task containing serial correlations: rats ignore the transition evidence accumulated across the trial history after error trials. This so-called reset strategy was highly robust and pervasive across many animals performing different task variants (Fig. 1). RNNs trained directly on the task did not replicate the reset strategy and were able to adopt the more optimal reverse strategy that leverages the binary symmetry of the 2AFC task (Fig. 2). On the other hand, RNNs pre-trained in an environment containing more than two alternatives exhibited the reset strategy displayed by the rats (Fig. 4). Furthermore, although the pre-trained RNNs did not show a transition bias after error trials, they maintained, as rats do, the transition history information and used it as soon as they made a correct choice (Fig. 5). Finally, we described a decoupling mechanism used by the pre-trained networks to gate off their transition bias after error trials (Fig. 6).

### Pre-training RNNs in ecologically relevant environments

RNNs have become a widely used tool to investigate the neural mechanisms underlying the behavior of animals in laboratory tasks (Mante et al. 2013; Sussillo 2014; Sussillo et al. 2015; Carnevale et al. 2015; Barak 2017; J. X. Wang et al. 2018; Remington et al. 2018; Mastrogiuseppe and Ostojic 2018; Yang and Wang 2021; Feulner and Clopath 2021; Saxena et al. 2021). The usual approach is to train the RNN directly on the task of interest and then investigate the underlying circuit mechanisms. By contrast, animals arrive at the laboratory with a plethora of priors that both guide and constrain the solutions learned by the animal to solve different tasks. By ignoring these priors RNNs are able to explore a much larger realm of strategies, frequently outperforming the animals but departing from the solutions they use. A quantitative discrepancy in performance has been frequently solved by adding noise to the units in the network (e.g. Mante et al., (Mante et al. 2013)) or to the stimulus (e.g. Sohn et al., (Sohn et al., n.d.)). However, although noise may be a mechanism limiting the performance of neural circuits (Faisal, Selen, and Wolpert 2008), it cannot account for all factors that constrain the space of solutions available to the animal. A complementary approach is to train the RNNs in several cognitive tasks, hence forcing them to develop strategies that are general enough to assure a good performance in all tasks (J. X. Wang et al. 2018; Yang et al. 2019; Hadsell et al. 2020). Here we followed a more specific approach (Ma and Peters 2020; Yang and Molano-Mazón 2021): pre-training the RNNs in a more naturalistic environment that can present more than two alternatives to address a qualitative discrepancy between the behavior of networks and rats in a 2AFC task with serial correlations.

This pre-training approach has been rarely used in neuroscience and, to the best of our knowledge, only for feedforward networks (Stoianov and Zorzi 2012; Roseboom et al. 2019). Nevertheless, within the machine learning (ML) community, pre-training is a routine procedure aiming to speed up the learning of a given task by assuring a good performance in a more general one (Dahl et al. 2012; Devlin et al. 2018; Tan and Le 2019). The structural priors induced by such pre-training help in naturalistic tasks like the ones used in ML, but can be a handicap in the tasks used in the laboratory that present oversimplified statistics or artificial symmetries. For instance, the visual system of humans possess extraordinary capabilities that are highly adapted to the statistics of natural scenes (Geisler 2008) but fail to correctly interpret seemingly simple stimuli, producing a visual percept that consistently differs from reality (i.e. a visual illusion) (e.g. (Weiss, Simoncelli, and Adelson 2002)). Similar phenomena have also been studied at a cognitive level (Kahneman 2011). For example, humans show a tendency to avoid losses at the expense of disregarding potential gains (Kahneman and Tversky 2012) a behavior that may have been useful in the past, helping to make fast decisions in dangerous environments (Kahneman 2011). Our results illustrate both sides of structural priors, with the NAFC pre-training helping the networks to leverage the transition history after correct responses but impeding them to do so after an error (Supp. Fig. 7c).

### Modeling the behavior of rats in the 2AFC task using RNNs

We chose RNNs to model the impact of structural priors on rats’ behavior because modern training methods allowed us to explore a vast realm of solutions in a relatively hypothesis-free fashion (i.e. without the choice of model introducing any obvious priors). Moreover, in contrast with more phenomenological models of behavior, RNNs offered us the possibility to characterize the neural mechanisms underlying the reset strategy (Fig. 6). The results from this analysis allowed us to formulate hypotheses and could guide the analysis and interpretation of rats’ neural recordings.

The pre-training protocol we used with the pre-trained RNNs cannot be but a simplification of the processes that the brain underwent during evolution and task learning. First, in our approach, the training of the RNN in the laboratory task was embedded in the pre-training in the NAFC environment (Fig. 3e). Importantly, the NAFC was not pre-designed to render the reset strategy as optimal: exploiting serial correlations in the trial sequence is always optimal, both after correct responses and after errors. Nevertheless, because the benefit of reversing after-errors decreases with N it is reasonable to expect that the networks will approach the reset strategy as N increases (e.g. N=14, 15, and 16). Because changes in N were blocked and cued to the RNN (i.e. by making zero all the stimulus dimensions higher than N) RNNs could have adapted their strategy to leverage the serial correlations on a block-by-block bases, just as they adapted their output choice behavior (they very rarely made choices for invalid dimensions above N; Supp. Fig. 15e). And yet, in this simplified NAFC environment RNNs resolved the challenge of being trained with different N by finding the same general strategy for all Ns: the reset. A potentially more realistic approach to approximate the multi-timescale learning of biological neural circuits, would be to build the connections of the RNNs using two different processes working at different time scales (Yang and Molano-Mazón 2021). For instance, a slow learning process similar to the one used here would simulate the effect of evolution, whereas a faster training, limited to certain connections and based on reward-modulated Hebbian plasticity (Miconi 2017), would simulate the learning of the laboratory task.

Second, we reduced the statistical structure of a “natural environment” to the NAFC setup containing the basic features needed to induce the transition bias: a decision task with multiple choices that presents varying statistical contexts, each characterized with a simplified structure in their transition probability matrices (Fig. 3d). Introducing more than three contexts or more complex transition matrices would have required longer training and possibly larger networks but we do not foresee an *a priori* limitation in this regard. A possible alternative to the statistics we used in the NAFC environment is to generate the state sequences independently (i.e. no serial correlations), with each context being defined by a different vector of heterogeneous alternative probabilities (Abbott et al. 2017). This pre-training would promote a structural prior adapted to estimate the first order rates of each alternative and a history bias which would tend to repeat rewarded options and stay away from unrewarded options. But it would not generate any gating of the accumulated evidence after errors. Interestingly, this type of history bias is commonly observed in the form of win-stay lose-switch bias in 2AFC tasks using uncorrelated sequences (Busse et al. 2011; Donahue, Seo, and Lee 2013; Akaishi et al. 2014; Frund, Wichmann, and Macke 2014; Abrahamyan et al. 2016; Urai et al. 2019) and in our animals in the 2AFC task with serial correlations (Hermoso-Mendizabal et al. 2020), raising the question of whether RNNs pre-trained in such statistics could indeed reproduce this ubiquitous behavior.

### Reinforcement versus supervised learning

RNNs are commonly trained with Supervised Learning (SL) protocols (Werbos 1990), which provide them with the correct answer at each timestep. Thus, networks not only know when they are wrong but also what they should have done. Here we have trained the networks using Reinforcement Learning techniques (RL) (Sutton and Barto 2018) which are primarily based on sparse feedback that reinforces the correct choices and discourages the mistakes, akin to the way animals learn in the laboratory (Niv 2009). This approach is a key feature of our training protocol, since the (lack of) access to the identity of the previous correct choice is at the core of the asymmetry between correct and error responses. Accordingly, when pre-trained networks had access to the previous correct answer they were able to adopt the more optimal reverse strategy (Supp. Fig. 7a). To the best of our knowledge, the extent to which SL and RL techniques promote different solutions for a given cognitive task has not been systematically explored. However, the differences at the behavioral (Lyon and Kuchling 2021) and neural (Donahue, Seo, and Lee 2013; Purcell and Kiani 2016a) level between after-error and after-correct responses should serve as a warning when comparing SL-trained networks with the activity of biological neural circuits.

### Neural mechanisms underlying the transition bias

To make informed decisions animals constantly integrate information at different temporal scales (Botvinick et al. 2019; J. X. Wang 2021). Yet, how this integration is implemented at the neural level is poorly understood (Schaeffer et al., 2021). Our population analyses of the activity of pre-trained RNNs provide an easy-to-use pipeline that can be directly applied to answer this question. Furthermore, they yielded specific predictions that can be tested in future experiments. First, our analyses showed that all variables involved in the computation of the transition bias were linearly decodable even before the stimulus is presented. Whether this is true in the brain of the rat remains to be fully tested. Several studies have suggested that representations in the brain are high dimensional, allowing for a simple linear readout of the variables of interest (Rigotti et al. 2013; Fusi, Miller, and Rigotti 2016). However, this might prevent the system from generalizing between similar scenarios (Stringer et al. 2019; Bernardi et al. 2020; Jazayeri and Ostojic 2021). In our paradigm, linear decodability is particularly surprising in the case of the encoding of the context (Fig. 6b), a latent variable that only impacts the choice of the network indirectly (combined with the previous choice) and thus could be implicitly encoded in a non-linear fashion. In the future, it will be important to elucidate the generality of this result by testing whether similar latent variables can always be decoded linearly from biological brains, where complex circuit architectures could in principle allow for different encoding-decoding schemes. Indeed, a possible extension of the present study is to investigate how trial history information is encoded in networks with more elaborate architectures (Yang and Molano-Mazón 2021) where other forms of communication within and between areas are available. For instance, feedback connections could allow for a representation in upstream areas that changes dynamically as a function of the inputs from downstream areas and thus cannot be linearly decoded. Second, our results show that pre-trained RNNs gate off the transition bias after error trials by decoupling the projection of the network activity along the transition bias axis from the final choice, while preserving the impact of the stimulus (Fig. 6h-i). Importantly, a transition bias, consistent with the reversal strategy, could still be decoded after errors (Fig. 6f-g). Thus the decoupling of the transition bias from the choice was the key mechanism implementing the after-error reset strategy, preserving the encoding of the context (Fig. 6b) and allowing the recovery of the transition bias after the next correct choice (Fig. 5a-c,e). Previous studies have shown that the brain uses private, orthogonal subspaces to disentangle information about different aspects of a task (Mante et al. 2013; Kaufman et al. 2014; Sheahan, Franklin, and Wolpert 2016; Bagur et al. 2018; Flesch et al. 2022). For example, population recordings in the premotor cortex during a context-dependent perceptual task show that monkeys use the context cue to “displace” the state of the network such that the encoding of the irrelevant stimulus feature is decoupled from the choice axis (Mante et al. 2013). Whether the brain of the rat uses the lack of reward as a cue to disregard the information about the transition bias, yet maintaining it in a private subspace, or whether it directly eliminates such information remains to be tested. On the one hand, areas of the brain such as the prefrontal cortex seem to keep task-irrelevant information while performing a given task, presumably to have access to it, in case it becomes useful in the future (Barraclough, Conroy, and Lee 2004; Xiong, Znamenskiy, and Zador 2015; Cazettes et al. 2021). On the other hand, representing information in the spiking activity of populations of neurons is costly. Therefore, in a scenario in which activity is expensive, the decoupling mechanism we found in the networks might be replaced by one that simply switches off the activity along the transition bias dimension. Alternatively, the disconnection could be done higher in the computation scheme (Fig. 6d) by for instance eliminating the encoding of the previous choice, something that would effectively prevent the computation of the bias (Fig. 6d). In summary, our population analyses highlight specific questions to explore when analyzing neural data recorded from animals performing a laboratory task that requires integrating information at different temporal scales (Abbott et al. 2017; Schaeffer et al., 2021; Sarafyazd and Jazayeri 2019).

### Processing feedback in correct versus error responses

Previous studies have also reported reset behaviors after erroneous choices. In a sensorimotor mapping task, monkeys trained to associate two visual stimuli with two different actions reverted to chance performance after making a single mistake (Fusi et al. 2007). This after-error reset of the stimulus-response association contrasts with the reverse behavior an optimal agent would develop in this task. Rats also showed a reset in their strategy, abandoning an exploitation strategy and adopting an exploratory behavior upon detection of a contingency change in a two-alternative task (Karlsson, Tervo, and Karpova 2012). In light of our findings, these two seemingly suboptimal behaviors may follow from the same principle: animals have adapted to environments with multiple alternatives where identifying the best exploitation strategy upon a contingency change may require some exploration.

The main hypothesis of the current work relies on a simple observation: in real life, rewarded actions reinforce the existing action policy (Sutton and Barto 2018), while errors are only informative about what not to do. This uncertainty after erroneous choices is magnified in scenarios that require decisions at different hierarchical levels beyond the perceptual categorization process: from the strategy to follow (Purcell and Kiani 2016b; Sarafyazd and Jazayeri 2019), to the exact motor trajectory (McDougle et al. 2016) or the timing to execute the selected action (Hernández-Navarro et al. 2021). This multi-level process makes the asymmetry between positive and negative outcomes larger. Such an asymmetry can be viewed as a direct consequence of the so-called Anna Karenina Principle, which states that there are many ways in which things can go wrong but only one in which they will go right (Diamond 2017). Hence, inferences about the environment after erroneous decisions are difficult because there are countless explanations about what went wrong. The extent to which all happy families are alike is an open question; our findings suggest, however, that our cognitive abilities can indeed be quite alike as a consequence of our shared evolutionary history in a highly structured world.

## Methods

All experimental procedures were approved by the local ethics committee (Comité d’Experimentació Animal, Universitat de Barcelona, Spain, Ref 390/14).

### Experimental methods

#### Animal Subjects

Animals were male Long-Evans rats (Charles River), pair-housed during behavioral training and kept in stable conditions of temperature (23 °C) and humidity (60%) with a constant light-dark cycle (12h:12h, experiments conducted during light phase). Rats had *ad libitum* food, and *ad libitum* water on days with no experimental sessions, but water was restricted during behavioral sessions.

#### Behavioral tasks

*Frequency discrimination task* (Supp. Fig. 13a): rats performed an auditory reaction-time two-alternative forced choice task (Hermoso-Mendizabal et al. 2020). Briefly, at each trial, an LED on the center port indicated that the rat could start the trial by poking in the center port. After a fixation period of 300 ms during which rats had to maintain their snout inside the port, the LED switched off and an acoustic stimulus consisting in a superposition of two amplitude-modulated frequencies was presented (see details below). Each frequency was associated with a specific side (high frequency left side; low frequency right side) and reward was available at one of the two lateral ports, depending on the dominant frequency. Animals could respond any time after the stimulus onset. Correct responses were rewarded with a 24 µL drop of water and incorrect responses were punished with a bright light and a 5 s time-out. Trials in which the rat did not make a side poke response within 4 seconds after leaving the center port were considered invalid trials and were excluded from the analysis (average of 0.4% invalid trials per animal). Withdrawal from the center port before stimulus onset (fixation break) canceled stimulus presentation. After a fixation break, rats were allowed to initiate fixation again.

*Intensity level discrimination task* (Pardo-Vazquez et al. 2019; Hermoso-Mendizabal et al. 2020) (Supp. Fig. 13b). Two speakers positioned at both sides of the box simultaneously played an amplitude-modulated white noise (see below for details). Rats had to discriminate the side with the loudest sound (right or left) and seek reward in the associated port. The rest of the details of the task are the same as in the frequency discrimination task.

*Silent task (Duque and de la Rocha 2022)* (Supp. Fig. 13c). At each trial, an LED on the center port acted as a go cue indicating the rat could start the trial by poking in. After a fixation period of 300 ms, the LED switched off and rats had to guess at which of the two lateral ports the reward would appear. The rest of the details of the task are the same as in the frequency discrimination task (Duque and de la Rocha 2022).

All the experiments were conducted in custom-made operant conditioning cages, the behavioral set-up was controlled by an Arduino-powered device (BPod v0.5, Sanworks, LLC, Stony Brook, NY, USA) and the task was run using the open-source software PyBPod (pybpod.com).

*Groups.* **Group Freq.**: frequency discrimination task with Prep = 0.7, n= 10 (Hermoso-Mendizabal et al. 2020). Intensity level discrimination (ILD) tasks: **Group ILD-0.8** with Prep = 0.8, n=8. **Group ILD-0.95** with Prep = 0.95, n=6. **Group ILD-Uncorr** with uncorrelated sequences (i.e. Prep= 0.5), n=6 (rats from Group ILD-0.8 before being trained with the correlated trials). Three of these rats did not develop any significant transition bias and were not used to compute the Reset Index. Silent tasks without any stimuli: **Group S0.8** with Prep = 0.8:, n=5. **Group S0.95** with Prep = 0.95: n=4.

|  | # rats | # sessions (mean, range) | # trials (mean, range) |
| --- | --- | --- | --- |
| FD task 0.7 | 10 | 59 | 56506 |
| ILD task 0.8 | 8 | 64 [21-87] | 39299 [11532-55792] |
| ILD task 0.95 | 6 | 16 [15-17] | 7666 [6523,8786] |
| ILD task Unc | 3 | 30 [28-34] | 18903 [14840-26516] |
| Silent task 0.8 | 5 | 19 [18-21] | 14380 [8940-19412] |
| Silent task 0.95 | 4 | 52 [44-60] | 31190 [18512-38307] |

#### Acoustic stimulus

In the two acoustic tasks used, the stimulus *S_k_(t)* was created by simultaneously playing two amplitude-modulated (AM) sounds *T_R_(t)* and *T_L_(t)*:

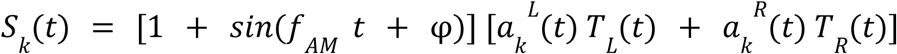

Where the frequency was *f*_AM_ = 20 Hz. The phase delay φ = 3π/2 made the envelope zero at *t* = 0. In the frequency discrimination task, *T_L_(t)* and *T_R_(t)* were pure tones with frequencies 6.5 kHz and 31 kHz, respectively, played simultaneously in the two speakers (Supp. Fig. 13e-h). In the interaural level discrimination task (ILD), they were broadband noise bursts played on the left and on the right speaker, respectively. The amplitudes of the sounds *T_L_(t)* and *T_R_(t)* were calibrated at 65 dB SPL using a free-field microphone (Med Associates Inc, ANL-940-1). Sounds were delivered through generic electromagnetic dynamic speakers ( ZT-026 YuXi) located on each side of the chamber.

#### Stimulus Sequence

A two-state Markov chain parametrized by the conditioned probabilities P_REP_ =P(L|L) and P(R|R) generated a sequence of stimulus category *c_k_* = {-1,1}, which determined the side of the reward (left/right) (Supp. Fig. 13d). In each trial the stimulus strength *s_k_* was randomly drawn from a fixed set of values = [0, 0.25, 0.5, 1]. *s_k_* defined the relative weights of the rewarded and non-rewarded sounds. The stimulus evidence was defined in each trial as the combination *e_k_* = *c_k_*\**s_k_*, thus generating seven different options (0, ±0.25, ±0.5, ±1) (Supp. Fig. 13). The value of *e_k_* determined the the p.d.f. from which the two sets of instantaneous evidences 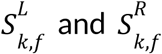 were drawn at each 50 ms frame *f* (Hermoso-Mendizabal et al., 2020). When *e* : = ±1 the p.d.f. for 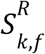 was *f*(*x*)=δ(*x*∓1) (i.e., a Dirac delta p.d.f.), whereas when *e_k_*∈ (−1,1), it was a stretched beta distribution with support [−1,1], mean equal to *e*_k_ and variance equal to 0.06. The p.d.f. Distribution for 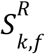 was the mirror image with respect to zero (Supp. Fig. 13). Finally, the amplitudes 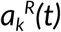 of the two AM envelopes were obtained using (Supp. Fig. 13):

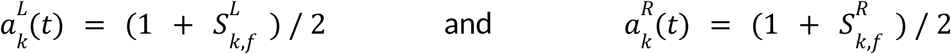

with *f* referring to the frame index that corresponds to the time *t*.

### Analysis of Behavioral Data

#### Psychometric curve analysis

The repeating psychometric curves were obtained for both rat experiments and RNN simulations, by computing the proportion of repeated responses as a function of the repeating stimulus evidence (*ê*) defined for the *t*-th trial as *ê_t_* = *r_t_*_-1_*e_t_*, with *r*_t−1_ = {−1,1}, representing the response in the previous trial (left or right, respectively). Thus, positive (negative) values of *ê* corresponded to trials in which the stimulus evidence pointed to repeating (alternating) the previous choice, respectively. Psychometric curves were fitted to a 2-parameter probit function:

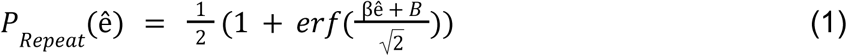

The sensitivity β quantified the stimulus discrimination ability, while the fixed side bias B captured the animal side preference for the left (B < 0) or right port (B > 0).

#### Generalized linear model (GLM)

For both rats and RNNs we fitted the same GLM model to quantify the weight that the various factors of the task had on the choices (Busse et al. 2011; Frund, Wichmann, and Macke 2014; Abrahamyan et al. 2016; Braun, Urai, and Donner 2018; Hermoso-Mendizabal et al. 2020) (Supp. Fig. 1). The probability *p*(*r_t_* =+ 1|*y_t_*) that the response *r_t_* at trial t was Rightwards was modeled as a linear combination of the current stimulus and trial history passed through the logistic function (Supp. Fig. 1e):

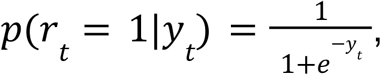

where the argument of the function in trial *t* was

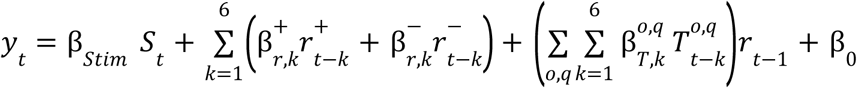

The current stimulus was given by *S_t_* defined as the average intensity difference between the two tone sounds in trial *t* (Supp. Fig. 1b). The trial history contributions included the impact of the previous ten trials (t-1, t-2, t-3…; grouping the impact of trials t−6 to trial t−10 in one term). The terms 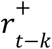 represented the previous rewarded responses being −1 (correct left), +1 (correct right), or 0 (error response). Similarly, 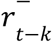 represented previous unrewarded responses being −1 (incorrect left), +1 (incorrect right), or 0 (correct response) (Supp. Fig. 1c). Previous transitions were given by the product of the last two responses:

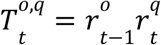

Meaning that they were 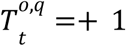 for repetitions and 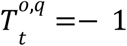 for alternations. The superindices {o,q} refer to the type of transition which depended on the outcomes of the last two trials *t*-1 and *t,* respectively: correct-correct {+, +}, correct-error {+, −}, error-correct {−, +}, and error-error {−, −} (Supp. Fig. 1d). Finally, the coefficient **β**_0_ represented a fixed side bias.

#### Reset Index

The Reset Index (RI) quantifies the extent to which an agent follows the reset strategy. It is computed as

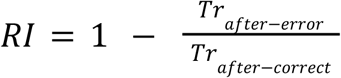

Where *Tr_after−error_* and *Tr_after−correct_* (Fig. 4g) are computed as the absolute value of the sum of transition weights 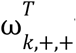 with k = 2, …, 6-10. Note that we excluded the most recent transition, k=1, from the computation to prevent lateral biases from affecting the index. The transition weights, 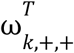, are obtained by separately fitting the GLM to after error and after correct trials, respectively. Because the Reset Index is only meaningful for agents that present a transition bias in the first place, we set a threshold (thcontr) and discarded any rat or network for which the sum of the transition history contributions (*Tr_after−error_* + *Tr_after−correct_*) was below the threshold. This only affected experiments with uncorrelated trial sequences (Group ILD-Uncorr, thcontr = 0.05) (Fig. 1g) and the analysis of the behavior of RNNs during training (Figs. 2e and 4f-g, thcontr=0.1). We obtained similar results for different values of thcontr.

### Simulations

#### Recurrent Neural Networks

All networks are fully connected, recurrent neural networks containing 1024 long short-term memory (LSTM) units (Hochreiter and Schmidhuber 1997). LSTM units are composed of an input gate, a forget gate and an output gate, each modulated by a learned function. This allows them to control which information enters the unit, which one is remembered and which one is output. The dynamics of the LSTM are defined by the following equations:

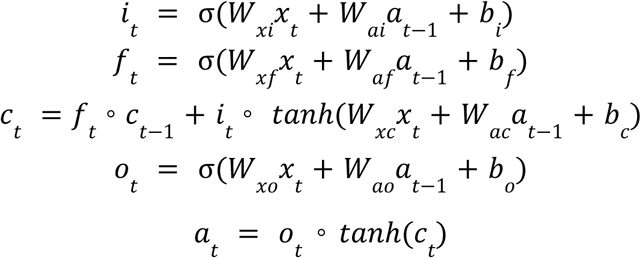

Where σ is the sigmoid function, and ∘ is an operator denoting element-wise multiplication; x_t_ is the input to the network at time t: a fixation cue (0 at the decision period and 1 otherwise), N inputs corresponding to the stimuli associated with each choice, and the reward and action at t-1; a_t_ is a vector containing the activity of all units in the network at time t; c_t_ represents the cell-states, which can be interpreted as the units’ internal states; i_t_ represents the input gates that control the information that enters the cell-states; f_t_ represents the forget gates that control the information from the cell-states at time t-1 that is preserved; o_t_ represents the output gates that control how much the cell-states of each unit affects its activity. All matrices, W, were modified during training (see below). The decision made by the network at time t was obtained by linearly projecting the activity of all units, a_t_, into a space with dimension equal to N_max_+1, applying a softmax function to the resulting vector and randomly sampling from the resulting probability mass function (Fig 2A right).

#### RNN Tasks

All tasks were implemented using the NeuroGym toolbox (Molano-Mazon et al. 2022). Tasks were organized into trials. For simplicity, trials were short and had the following structure: a fixation period (lasting one timestep), stimulus (composed of either one or two timesteps, one sample per timestep, mean=1.3 samples) and decision period (one timestep). There was no inter-trial interval. At the decision period, the network had to choose the action associated with the stimulus that, averaged over the timesteps, was the largest. The reward given to the network was 0 throughout the trial, except at the decision timestep, when a reward of 1 was given to the network if it made the correct choice and 0 otherwise. To make sure that the short length of the trials employed did not affect our results, we ran simulations with longer trials: the fixation period lasted 3 timesteps; stimulus duration was on average 4 timesteps (min=2, max=6 samples) and the decision period was 3 timesteps. We confirmed that when trained using these longer trials, networks could learn the task and show the same reset behavior (Supp. Fig. 14).

*Two-Alternative Forced Choice (2AFC) task*. As in the experiments with rats, the trial sequence of stimulus categories in the 2AFC task was generated by a two-state Markov chain parametrized by the repeating probability, Prep, which changed in trial blocks between Prep=0.8 (Repeating block) and Prep=0.2 (Alternating block) (Fig. 1a). At the end of every trial, the block could change with probability p = 0.0625, yielding an average block length of 160 trials. This length is comparable to the one used in the experiments (200 trials per block in group FD; 80 trials per block in groups ILD-0.8, ILD-0.95, S0.8, S0.95).

*N-Alter*native Forced Choice (NAFC) task pre-training (Supp. Fig. 15a-d). As in the 2AFC task, the trial sequence of stimulus categories is organized into blocks with distinct serial correlations. We pre-trained the networks using 3 different contexts (i.e. trial blocks): repeating, clockwise and anticlockwise (Fig. 3c-d). In each context, the sequences were generated by an N-state Markov chain parametrized by a different transition matrix (Fig. 3d). The transition matrix Pr(j|i) determined the probability of moving from category i to category j in the next trial (*i*, *j* = 1, 2, …, *N*). In the repeating context, all the entries in the main diagonal were equal to Pr(i|i)=0.8 and the rest of the terms were Pr(j|i)=0.2/(N-1) (when *i* ≠ *j*). For the clockwise context, the matrix was Pr(i+1|i)=0.8 and assumed a circular topology of the states meaning that from the last state there was a likely transition to the first state, (i.e. Pr(1|N)=0.8). The rest of the terms equal to 0.2/(N-1). In the anticlockwise context it was similar except that the high probability transitions were Pr(i-1|i)=0.8 and Pr(N|1)=0.8. At the end of every trial, the block could change with probability p = 0.008. Every 5000 trials, the number of available choices *N* was randomly selected between 2 and the maximum number of choices (Nmax). Except where specified (e.g. Supp. Fig. 7b-c), the percentage of trials with *N* = 2 was prefixed to 1% to assure that all pre-trained networks saw the same amount of 2AFC trials regardless of the maximum number of alternatives, N_max_, with which the network was being pre-trained.

*Stimuli.* After having generated the trial sequence of stimulus categories, we generated for each category, random stimuli with different values of the mean stimulus strength *s*. The stimuli were always composed of 1 or 2 temporal samples (except in Supp. Fig. 14 where they were composed of 2-6 samples) presented sequentially (see stimulus examples in Fig. 2a). For each sample, the stimuli were drawn from a multivariate normal distribution with dimension *N* equal to the number of choices. For each stimulus category *c* =1, 2, …N, we randomly and parametrically varied the mean stimulus strength *s* on a trial by trial basis. We used the following six values: *s* ∈ [0 , 0. 12, 0. 25, 0. 5, 1]. The value of *s* determined the mean µ = (µ_1_ , µ_2_ ,…, µ*_N_*) of the multivariate normal distribution. For the dimension corresponding to the category, the mean was µ*_i_*= 0. 5 + κ *s* (for *i* = *N*) whereas for the rest of the dimensions it was µ*_i_* = 0. 5 − κ *s* (for *i* ≠ *N*). The prefactor κ = 0. 92 *log*(*N*) − 0. 02 was used to compensate for the increase in difficulty associated with a larger number of choices. The multivariate distribution had a diagonal covariance matrix (samples were independent between dimensions) with a uniform standard deviation equal to σ = 0. 1 for all stimuli and contexts. In the case of the 2AFC task, we set the categories Left and Right as *c* = {-1,1}, respectively, and we defined the mean stimulus evidence as the product *e* = *c* × *s*. As done for the experimental data, we computed psychometric curves representing the probability of the network to repeat the previous choice as a function of the repeating stimulus evidence (*ê*) defined for the *t*-th trial as *ê_t_* = *r_t_*_-1_*e_t_*, with the previous response *r*_t−1_ = {−1,1} representing left or right, respectively (Figs. 2d, 4c; Eq. 1). In the NAFC environment, the number of choices *N* varied in 5000 trial blocks between 2 and *N_max_.* The dimensionality of the input stimuli was however kept constant at the maximum Nmax during the entire training but the stimuli were set to zero in the invalid dimensions *N* < *n* ≤ *N_max_* (Supp. Fig. 15a-d). This choice made it easy for the networks to identify the eligible choices *n* ≤ *N* as those in which the input was non-zero. The dimension of the network’s output was also kept constant at Nmax. For that reason, networks could always choose among the Nmax choices even if the number of valid choices N was *N* < *N_max_* . When the network’s chose an invalid category or did not choose (i.e. fixated during the decision period), the trial was counted as an error trial. On average, only a small fraction of trials were of this type (Supp. Fig. 15e). Apart from the input stimuli, networks received as input two scalars, indicating the reward received in the preceding timestep and the choice made in the preceding timestep (J. X. Wang et al. 2018) (Fig 2a left).

#### Training of the RNNs

Networks were (pre-) trained for ∼3M trials on the corresponding task (the 2AFC or the NAFC environment). The training was done using Reinforcement Learning (RL). Importantly, RL differs from standard Supervised Learning methods in that the network never has explicit access to the correct answer, and only receives a scalar (the reward) as feedback on its performance. The amount of reward accumulated across timesteps is used to compute a loss function to update the weights of the network using backpropagation through time (BPTT) (Werbos 1990). We used an actor-critic deep reinforcement learning algorithm with experience replay (ACER) (Z. Wang et al. 2016) (Stable-Baselines toolbox (https://github.com/hill-a/stable-baselines). Actor-critic methods use a separate function (the actor) to explicitly represent the policy, independently of the function (the critic, or value function) that represents the value of each environment state (Sutton and Barto 2018). Experience replay constitutes a biologically inspired mechanism (McClelland, McNaughton, and O’Reilly 1995; O’Neill et al. 2010) that allows the network to store its previous experiences and learn from them *off-line* after randomizing over the stored data, which removes correlations in the observation sequence (Lin 1992). We tested alternative algorithms (A2C (Mnih et al. 2016), ACKTR (Wu et al. 2017) and PPO2 (Schulman et al. 2017)) and chose ACER because it was the one that learned the NAFC environment faster. However, preliminary analyses of the networks trained with these different algorithms did not show any clear difference in the strategy they used to solve the task (data not shown).

*Training hyperparameters.* The learning rate was set to 7 × 10^−4^. The discount factor, which controls the importance that future rewards have on the loss function that is used to update the weights of the network, was set to 0.99. Weights were optimized using RMSProp. When training RNNs with BPTT, one needs to decide how many steps to propagate the gradients back in the past in order to update the weights. We set this hyperparameter to 15 timesteps (∼5 trials) in all experiments except for the ones shown in Supp. Fig. 14, where the rollout was 50 (∼6 trials) to account for the longer duration of trials. ACER allows parallely acquiring experience via different agents (threads) and using those experiences to update the weights of the network. In all our experiments, the number of threads was set to 20. Other hyperparameters were kept as in the original paper (Z. Wang et al. 2016): weight for the loss on the Q value = 0.5, weight for the entropy loss = 1*i*10^−2^, clipping value for the maximum gradient = 10, scheduler for the learning rate update = ’linear’, RMSProp decay parameter = 0.99, RMSProp epsilon = 1 × 10^−5^ , buffer size in number of steps = 5 × 10^3^ , number of replay learning per on policy learning on average, using a poisson distribution = 4, minimum number of steps in the buffer, before learning replay =1 × 1 ^3^ importance weight clipping factor = 10.0, decay rate for the Exponential moving average of the parameters = 0.99, max KL divergence between the old policy and updated policy = 1.

#### Testing the RNNs

After the training, the networks were tested on the standard 2AFC task, with their weights frozen. Therefore, during the testing, networks can adapt their choice biases to the different blocks by changing their neural activity but not their connection weights.

#### Perfect integrator agent

The perfect integrator agent in Fig. 2b-c is simulated by summing all the stimulus samples presented in a given trial and selecting the choice associated with the largest resulting value.

### Analysis of Network activity

#### Linear Support Vector Machines (SVM)

We trained linear support vector machines (SVM) in (1) networks pre-trained on the N-AFC task (pre-trained networks; n=16), and (2) in networks directly trained on the 2AFC task (2AFC-trained networks; n= 16). The SVMs were designed for separating the following binary categories: (1) the local context (repeating versus alternating), (2) upcoming stimulus category (left versus right) and (3) the previous network’s choice (left versus right). We used the network activity, a_t_, of all the 1024 units in each RNN from the *training dataset* (see below) and the ground truth binary categories as the corresponding labels. We used the neural activity during the testing on the 2AFC task, i.e. while the network weights were frozen, taken from the fixation period, i.e. right before the stimulus of the current trial was presented. Once the SVM was trained, we obtained the maximal marginal hyperplane separating the two categories and the normal vector to the hyperplane which was defined as the decoding axis (Fig. 6a). The projection of the neural activity on this axis yielded a scalar coordinate that quantified the magnitude of the network encoding for the quantity of interest. Because we trained one SVM to predict the current stimulus category before the stimulus was actually presented, we interpreted this prediction as a proxy of the transition bias.

#### Decoder training methods

Trials were first labeled as after-correct (AC) and after-error (AE) trials according to the outcome of the last choice *r_t_*_−1_ Among all trials, the *training dataset* was made of trials, both AC and AE, with *clear context,* meaning that they were preceded by a clear repeating or alternating sequence of choices. The rationale of this choice was to facilitate the training by using trials in which the RNN could have a clear representation of context because it had just followed a consistent pattern of transitions. Clear context trials were defined as trials preceded by a sequence of choices *r_t_*_−*k*_ which were correct for *k* = 2, 3 and 4, and which were either all repetitions (*r_t−k−1_* = *r_t−k_*) or all alternations (*r_t−k−1_* ≠ *r_t−k_*). For each network, we trained the decoders using 500 clear context trials (half AE and half AC; Fig. 6). Once trained, we tested the accuracy of the encoders by separately projecting either 500 AC trials or AE trials drawn randomly from all trials (Fig. 6b, g-i). We repeated this procedure 500 times for each RNN and obtained the mean value (correlation). Our measures of encoding accuracy (see below) were thus cross-validated because of the separation between the training dataset (clear-context trials, ∼32% of total trials) and testing dataset (all trials). For visualization purposes, we used the encoding coordinates obtained from clear context trials only (3D and 2D scatter plots in Fig. 6e-f)

To control that differences in the transition bias encoding between AC and AE trials were not caused by training the decoders with a mix of AC and AE trials, we independently trained different decoders using only AC or AE trials from the training dataset (i.e. with clear context). Hence we obtained two separate decoding axes for each category. We then tested each decoder using trials with the same previous outcome randomly selected from all trials (e.g. we used AC trials to test the decoder trained using AC trials; Supp. Fig. 12).

#### Context Encoding accuracy

To quantify the accuracy of the context encoding we obtained the trial-by-trial encoding by projecting the activity of all trials (∼25K AC trials and ∼6K AE) on the Context axis (Fig. 6b). Using the true block label of each trial, we obtained two histograms (Fig. 6b right). We then used the Area Under the Receiver Operating Characteristics curve (AUC-ROC) as a measure of how well the current block (i.e. repeating or alternating) could be inferred from the neural encoding of the context (Fawcett 2006). In other words, the AUC-ROC quantified how separable were the histograms of the context encoding shown in Fig. 6b. If AUC=0.5 the two distributions would be indistinguishable whereas when AUC=1 they would be perfectly separable.

#### Transition bias Accuracy

We randomly selected either 500 AC trials or 500 AE trials from *all trials* (i.e. without imposing a clear context) and obtained the trial-by-trial transition bias encoding by projecting their activity on the decoding axis for Transition bias. We did this for each of the 500 training instantiations of the transition decoder. To quantify the accuracy of the transition bias encoding in predicting the upcoming stimulus category, we categorized the value of the bias into either Left or Right using a categorization threshold at zero. Accuracy was then obtained as the fraction of trials in which this categorical transition bias matched the true upcoming stimulus category (Fig. 6g). Notice that, because of the stochastic design of the task, this accuracy cannot be larger than 0.8. This is because even in the case of a perfect agent that has access in each trial to the context identity, the prediction about the upcoming stimulus will fail in the 20% incongruent trials, i.e. trials in which the transition was against the context tendency (e.g. a repetition in an alternating block).

#### Relationship between Context Encoding and Transition Bias

To quantify the relationship between the encoding of *context* and *transition bias*, and how it depended on previous outcome, we used the Pearson correlation coefficient between the two variables. The transition bias depends on context inversely depending on whether the previous choice was Left or Right. Because of the symmetry of the problem, we only used after Left choice trials for this analysis. We finally quantified this relationship separately for after-correct and after-error trials (Supp. Fig. 11). We assumed the arbitrary convention that positive context encoding corresponds to Alternation, and positive transition bias to Rightwards choice (Fig. 6d-e). Following this convention, positive values of the correlation coefficient mean that the more Alternating is the context encoding, the more towards the Right is the transition bias. This is the optimal thing to do after a Left correct choice (left panels in Supp. Fig. 11a-b). In contrast, after a Left error trial, the optimal thing to do would be to bias choices towards the Left in the Alternating context, implying a negative correlation between transition bias and context (right panels in Supp. Fig. 11a-b).

#### Impact of the Transition bias encoding on choice behavior

To quantify the impact of the transition bias encoding on the network’s behavior, we randomly selected 500 trials (either AC or AE; no constraints on previous transitions) for each of the 500 instantiations of the trained transition bias decoder, computed the transition bias in each trial (total num. trials 500 x 500), and binned the obtained values in four bins (range median ± 3 standard deviations). We then computed, for each bin and each mean stimulus evidence *e*, the fraction of trials in which the network made a rightwards choice (Fig. 6h). We made these psychometric functions separately for after-correct and after-error trials. To quantify the impact of transition bias on choice, we computed the slope of this function for zero stimulus evidence (Fig. 6i).

## Data code availability

Data and code will be made available on a public repository once the manuscript is published.

## Acknowledgements

We thank Jorge del Pozo for preliminary analyses. We thank Ainhoa Hermoso-Mendizabal for useful discussion and Lorena Jiménez for help with training of the animals. We thank the Barcelona Supercomputing Center (BSC) for providing computing resources. This research was supported by the Beatriu de Pinós fellowship, Generalitat de Catalunya (2017-BP-00305 to M.M.M), the Spanish Ministry of Economy and Competitiveness together with the European Regional Development Fund (IJCI-2016-29358 to D.D.; RTI2018-099750-B-I00 to J.R.), the European Research Council (ERC-2015-CoG - 683209 Priors to J.R. ) and the Simons Foundation Junior Fellowship, NSF NeuroNex Award 589 DBI-1707398, and the Gatsby Charitable Foundation which supported G.R.Y.. YS and SO were supported by the program “Ecoles Universitaires de Recherche” implemented by the French Agence Nationale de la Recherche under reference ANR-17-EURE-0017.

## Author contributions

M.M.M. and J.R. conceived the project; M.M.M., G.R.Y. and J.R. designed the model; D.D. and J.R. designed the experiments; D.D. carried the experiments; M.M.M. and D.D., analyzed the experimental data; M.M.M. trained and analyzed the behavior of the networks; M.M.M., D.D., G.R.Y. and J.R. interpreted the behavioral data; Y.S. and S.O. performed the population analyses; Y.S., S.O., M.M.M. and J.R. interpreted the population analyses; M.M.M. and J.R. wrote the manuscript with contributions from D.D., Y.S., G.R.Y. and S.O.

## Supplementary figures

**Supplementary figure 1.**
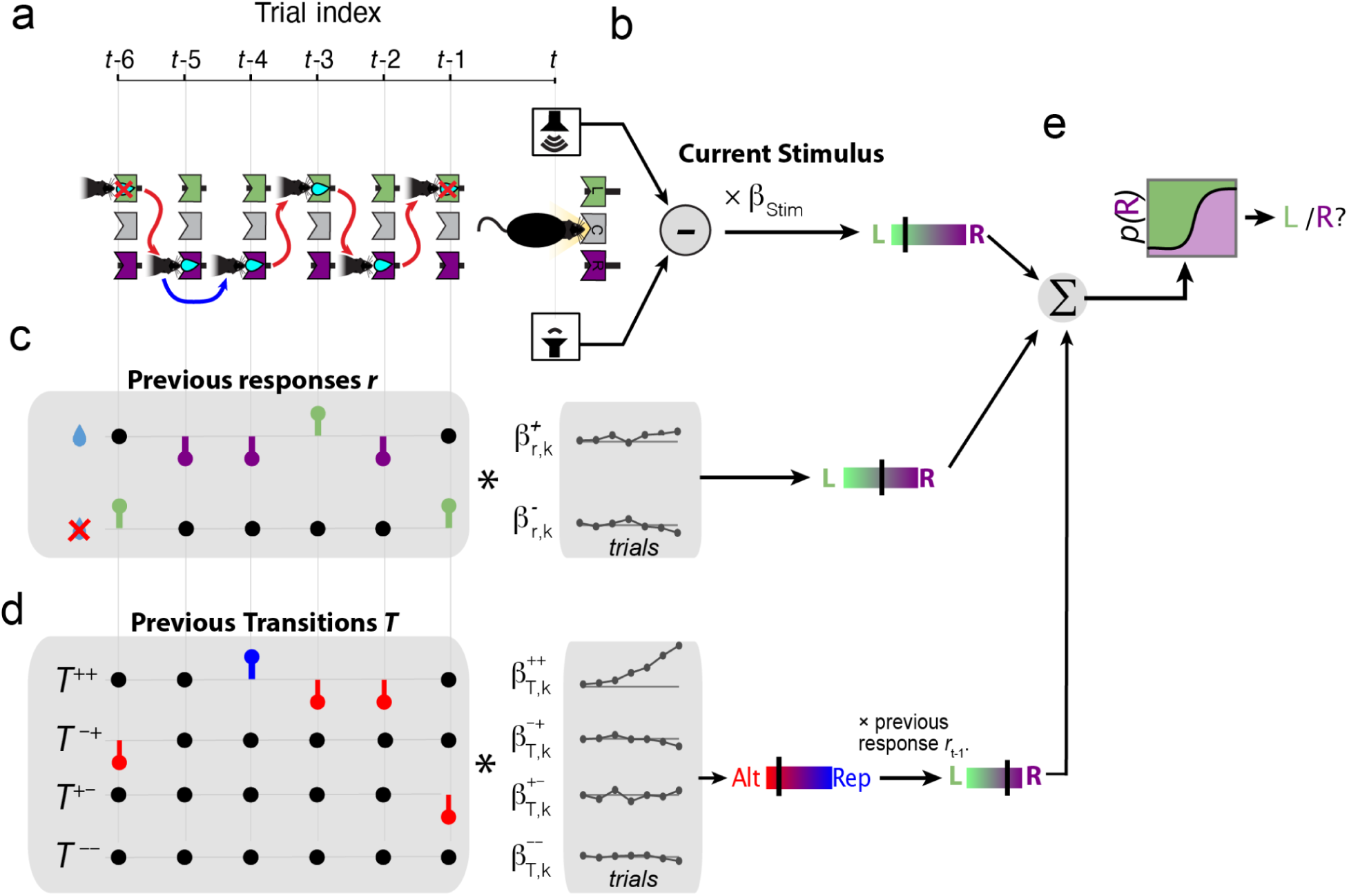
GLM assessing the impact of current stimulus and recent history onto rats’ choices. **a,** Exemplar series of recent history trials used to model rat decisions at current trial *t*. The Response series shows the rat decisions in the last six trials and the Transitions defined from the relation of two consecutive responses (blue arrow for Repetition, red arrow for Alternation). Outcome in each trial is indicated by a drop of water (correct) or a crossed-drop (error). These series are combined to generate the regressors for correct and error responses (c) and the four types of Transitions (d). **b,** The amplitude difference between the Left and the Right stimulus of the current stimulus is weighted by a single *β*_Stim_ . The outcome of this product provides the net evidence which can take values ranging from strong Left to strong Right (color bar with green-purple gradient; black tick shows the value of the example stimulus). **c,** The previous responses are weighted separately in rewarded *r*+ and error responses *r*⁻ that take the values -1 (Left), +1 (Right) or 0. Each of these two series convolved with the corresponding response kernels *β*^+^_r,k_ and *β*^-^_r,k_, respectively. **e,** Transitions are considered separately depending on the outcome of the two trials in the transition: *T*++ (rewarded-rewarded), *T*⁻+ (error-rewarded), *T*+⁻ (rewarded-error) and *T*⁻⁻ (error-error) which take the values -1 (alternation), +1 (repetition) and 0. The weighted sum of transition regressors is then multiplied by the previous response *r_t-1_* in order to yield the transition bias which provides Rightward vs Leftward evidence. **e,** The weighted sum of the stimulus, previous responses and previous transitions is then passed through a sigmoid function to yield the trial-by-trial probability of selecting a Right response. Individually for each rat or each RNN, the weights of the kernels (shown as black connected dots in the center gray boxes) were fitted to maximize the model’s probability to generate the actual choices.

**Supplementary figure 2.**
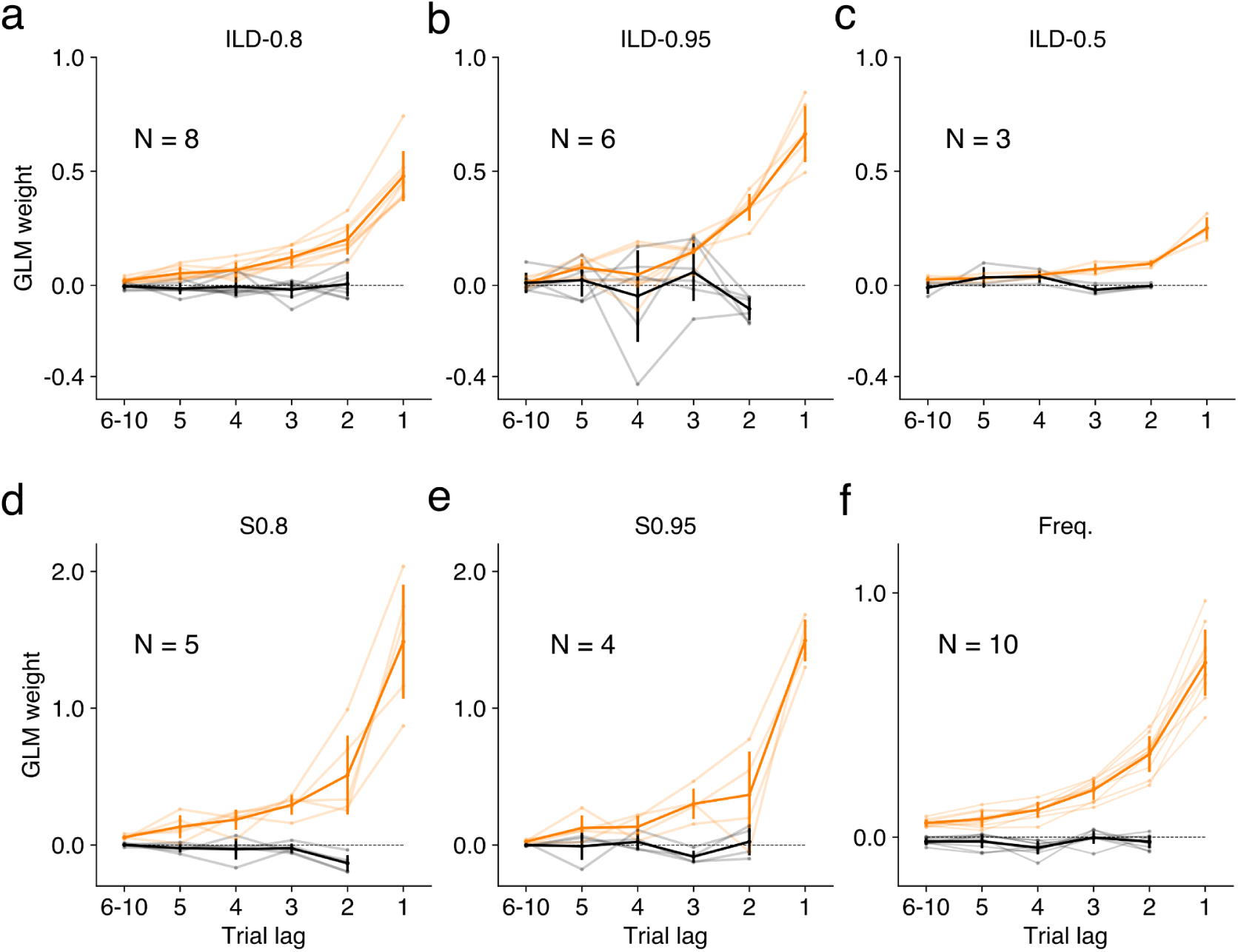
GLM weights of previous correct transitions (i.e. formed by two correct choices) computed after correct (orange) and after error (black) trials; mean and std. dev. (thick traces) and individual animals (light traces) from **a)** Group ILD-0.8: intensity level discrimination task with P_REP_ = 0.8, n=8 **b)** Group ILD-0.95: ILD task with P_REP_ = 0.95, n=6; **c)** Group ILD-Uncorr: ILD task with uncorrelated sequences, (i.e. P_REP_ = 0.5), n=3; **d)** Group S0.8: silent task (without any stimuli) and P_REP_ = 0.8, n=5; **e)** Group S0.95: silent task with P_REP_ = 0.95, n=4. **f)** Group Freq.: frequency discrimination task, n=10 (Hermoso-Mendizabal et al. 2020).

**Supplementary figure 3.**
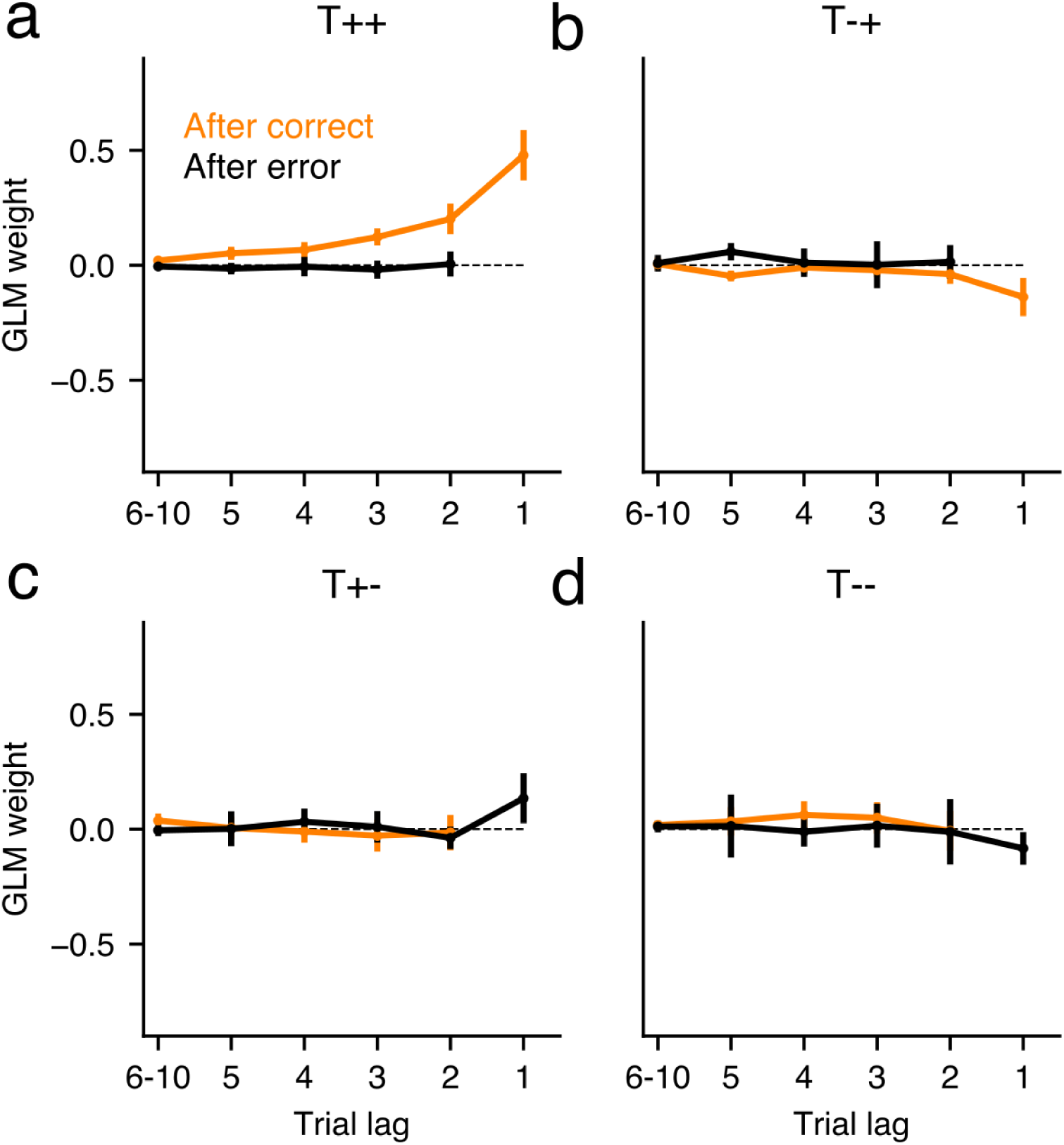
Only transitions composed by two correct choices influence the decision of rats. The figure shows all transition kernels obtained from the GLM separately fitted to after-correct and after-error trials (see color code in a). Each panel shows the contribution of the different types of transitions: T++: transitions made of two consecutive rewarded trials or correct-correct transitions **(a)**, T-+: error-correct transitions **(b)**, T+-: correct-error transitions **(c)**, T--: error-error transitions **(d)**. Dark lines show mean values (Group ILD-0.8, n=8). Error-bars correspond to standard deviation.

**Supplementary figure 4.**
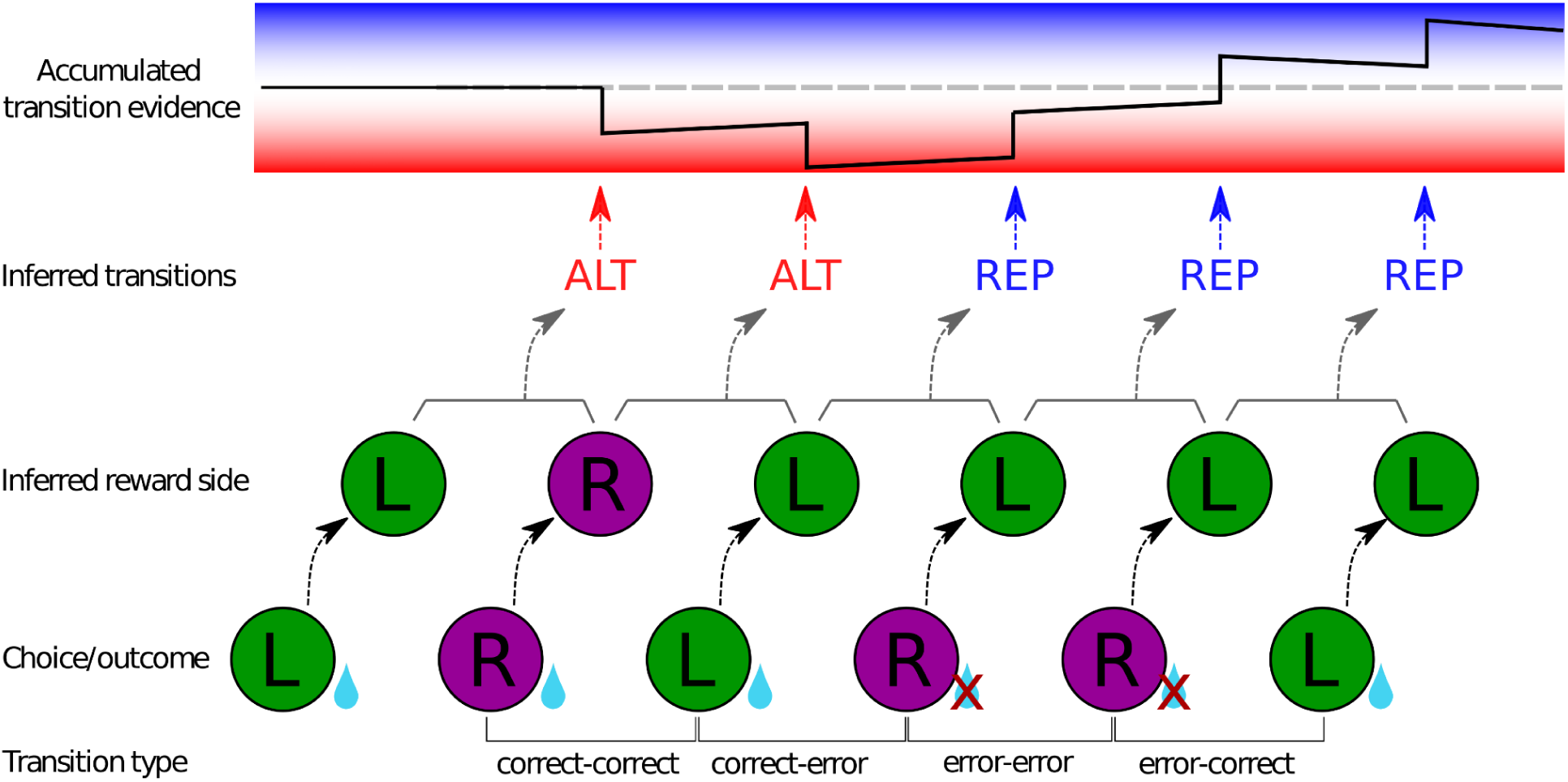
Inference of ground-truth reward sides and transitions, and accumulation of transition evidence made by an optimal agent for different transition types. Bottom: example sequence of choices and outcomes. The type of transition connecting each pair of consecutive choices is indicated in the bottom. **Middle:** after making a choice and learning the outcome, the agent infers the rewarded side (black dashed arrows). Having inferred two consecutive rewarded sides, the agent infers the last ground-truth transition (dark gray dashed arrows). **Top:** Each inferred ground-truth transition is used to update the accumulated transition evidence (blue/red dashed arrows). Notice that following this inference procedure, an optimal agent can use each transition to update the accumulated evidence independently of its type (i.e. ++, +-, -- and -+). The agent can then leverage this accumulated evidence to bias the upcoming choices (not shown for simplicity).

**Supplementary figure 5.**
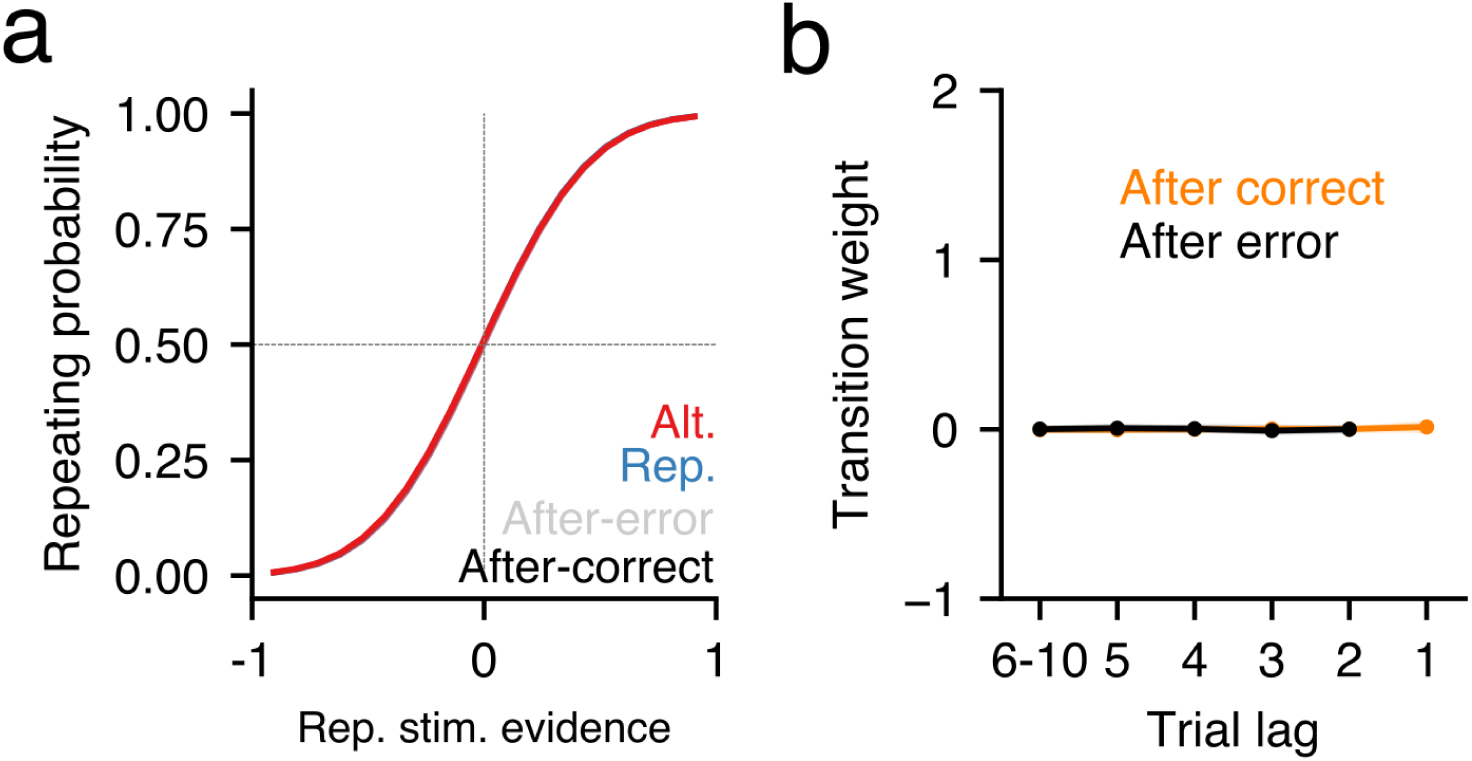
Networks trained in a 2AFC without trial-to-trial correlations. do not develop any transition bias. **a)** Psychometric curves obtained for each block (see color code) when testing an RNN trained on a standard 2AFC task without trial-to-trial correlations on the 2AFC task, computed in after-correct (dark) and after-error (light) trials. **b)** GLM weights of previous correct transitions (i.e. formed by two correct choices) computed after correct (orange) and after error (black) trials (n=8, thick traces) and individual networks (light traces).

**Supplementary figure 6.**
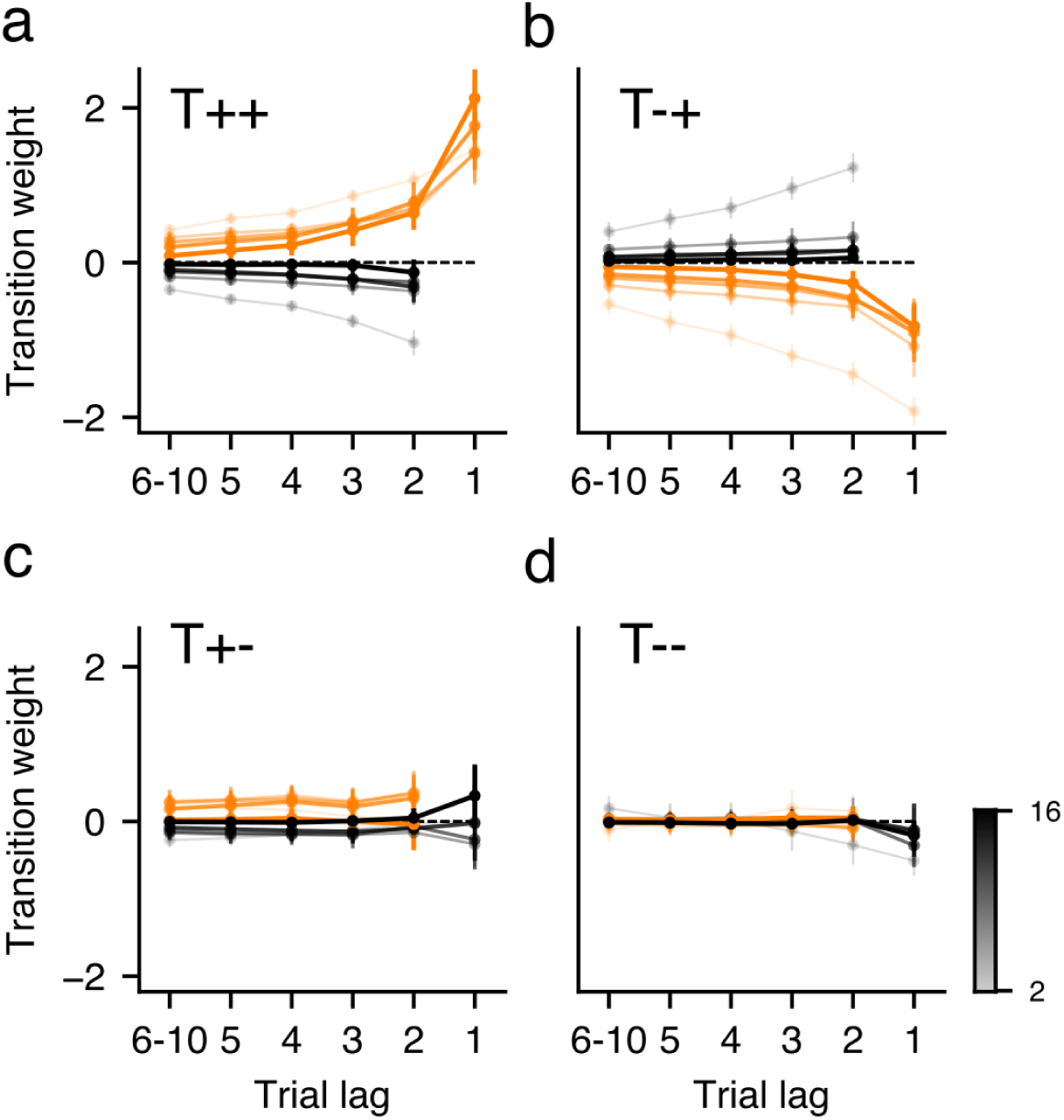
Average transition kernels obtained from pre-trained RNNs when testing in the 2AFC task, for pre-training NAFC environments with a different maximum number of alternatives, from N_max_=2 (thick, light traces) to N_max_=16 (thin, dark traces). Notice that as N_max_ increases, all kernels vanish , except the T++ after-correct choices.

**Supplementary figure 7.**
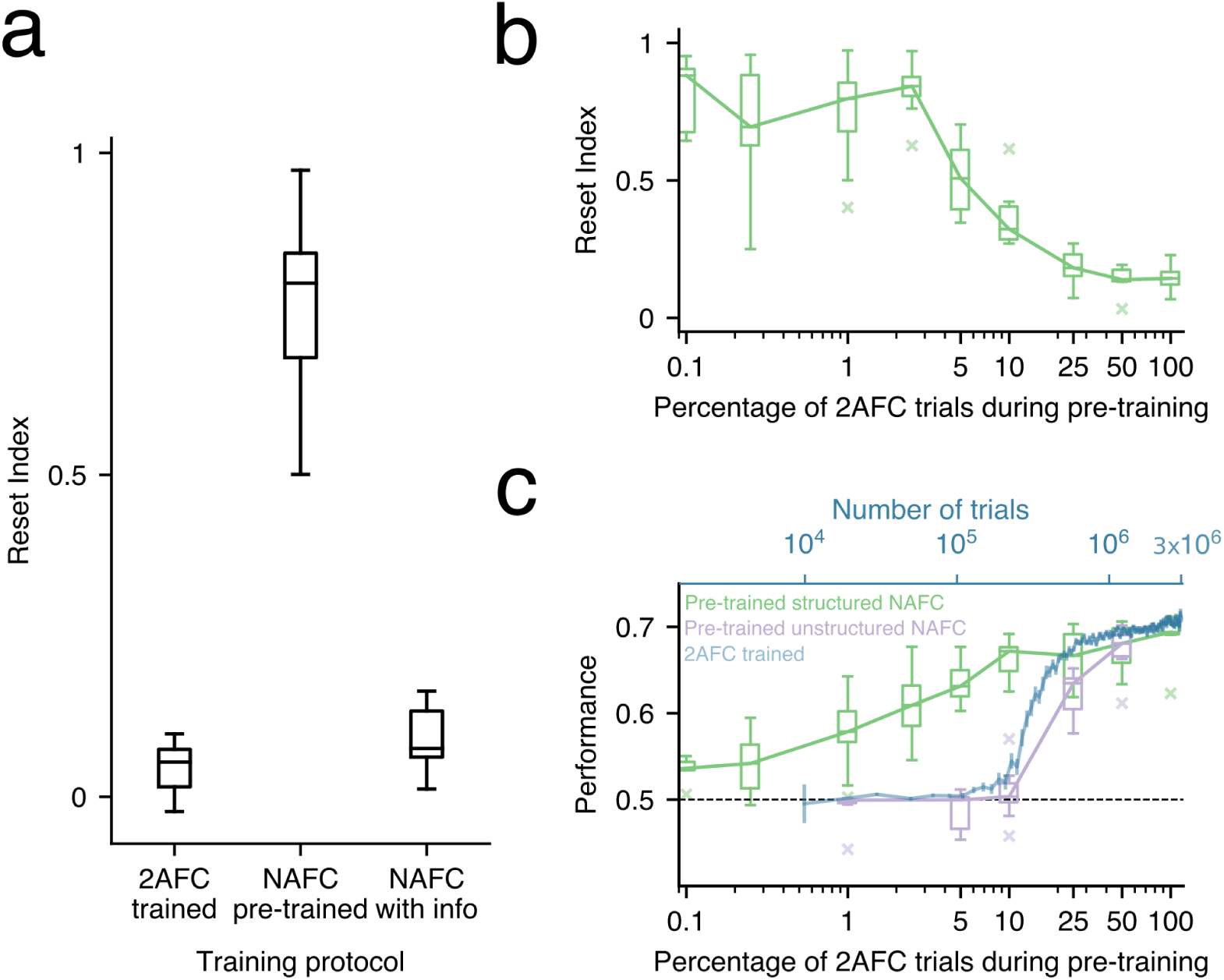
Reset strategy in pre-trained networks depends on features of pre-training. **a)** Comparison of the Reset index for RNNs directly trained in the 2AFC task (as in Fig. 2), pre-trained in standard NAFC (as in Fig. 4) and RNNs pre-trained in a variant of the NAFC environment in which we provided as an input to the network the correct alternative in the previous timestep (“NAFC with info”). **b-c)** Reset index (b) and accuracy (c) in the 2AFC task as a function of percentage of 2AFC trials embedded in the NAFC pre-training, for networks pre-trained in the standard NAFC (Structured NAFC, green) and networks pre-trained in a NAFC environment with no acros-trial correlations when N>2 but standar repeating and alternating blocks when N=2 (unstructured NAFC, purple). Only trials with no stimulus evidence were used to compute accuracy (c). As a reference, the performance of networks trained directly on the 2AFC task for a comparable amount of trials is shown (blue trace, top axis in c). Only RNNs that developed a transition bias were used to compute the Reset Index values. Both the structured and the unstructured NAFC pre-training were done with N_max_ = 16. x-axis in log-scale in b and c.

**Supplementary figure 8.**
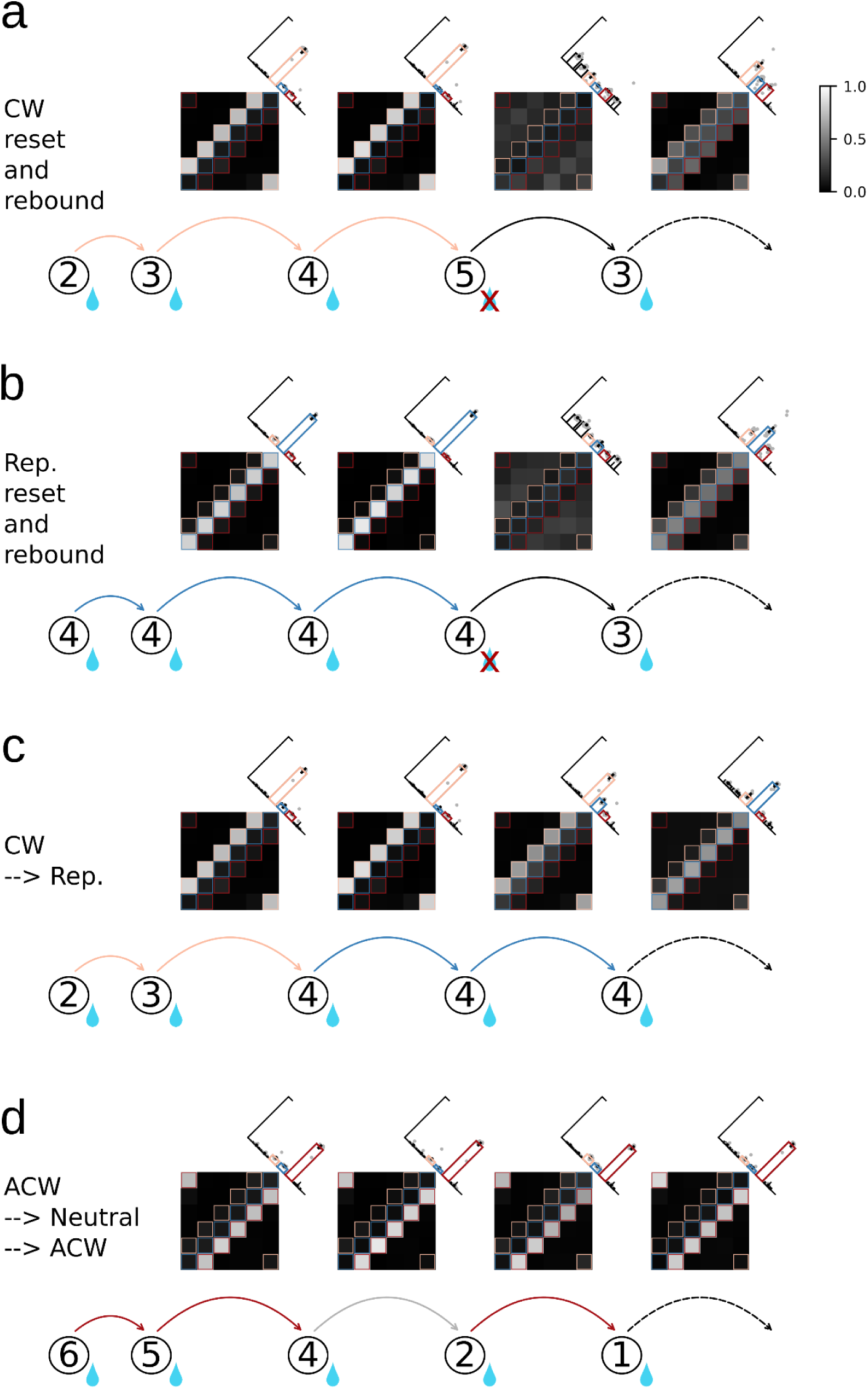
Evolution of the transition bias matrix computed using pre-trained RNNs (Nmax= 16, RNNs are tested in N=6) throughout different choice sequences. Insets above each matrix show histograms quantifying the trial-by-trial estimates of being in a Repeating (blue bar), Clockwise (pink) and Anticlockwise (red) context (as in Fig. 5). **a-b)** Evolution of the transition bias matrix throughout a choice sequence containing an error (as in Fig. 5) after a choice pattern consistent with the CW (a) and the Rep (b) contexts. **c)** Evolution of the transition bias matrix throughout a choice sequence involving an unexpected correct response occurring in context change-point (switching from the CW to the Rep context). The first unexpected correct transition at the change-point induces a decrease in the networks estimate to be in the old context and a slight increase in favor of the new context; the second unexpected correct transition makes the networks completely switch their internal estimate towards the new context. **d)** Evolution of the transition matrix throughout a sequence with correct unexpected neutral transition, i.e. a transition which is incongruent with any of the contexts. Neutral transitions had only a slight effect on the transition matrices. Note that this type of transitions constitute a significant portion of incongruent transitions which are themselves 20% of the total.

**Supplementary figure 9.**
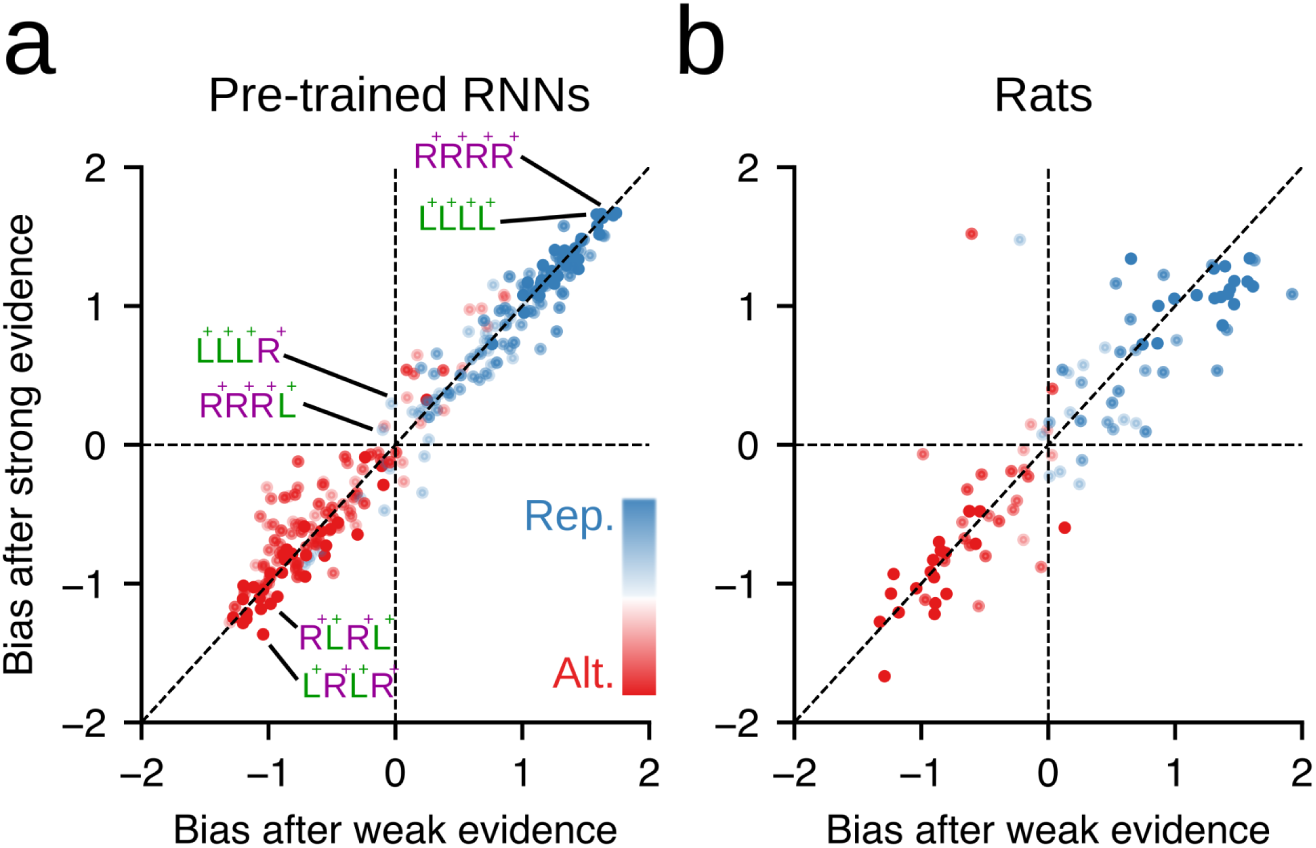
Transition bias does not depend on the magnitude of the stimulus evidence of the previous trial. **a-b)** Transition bias of pre-trained RNNs (a) and rats (b) computed after strong and after weak evidence stimuli. To isolate the impact of the previous stimulus evidence, we conditioned on the sequence of previous correct choices (total of 16 different 4-choice sequences; see insets pointing to example sequences). Each point represents the Transition bias for one sequence obtained from one agent (n=16 RNNs in a; n=10 rats from the Group Freq. in b) Stimuli were classified into strong and weak evidence using a median split of the stimulus strength *s*. Dot color represents the estimated transition bias obtained from convolving the transition kernels with the pattern of transitions of each sequence (see colorbar).

**Supplementary figure 10.**
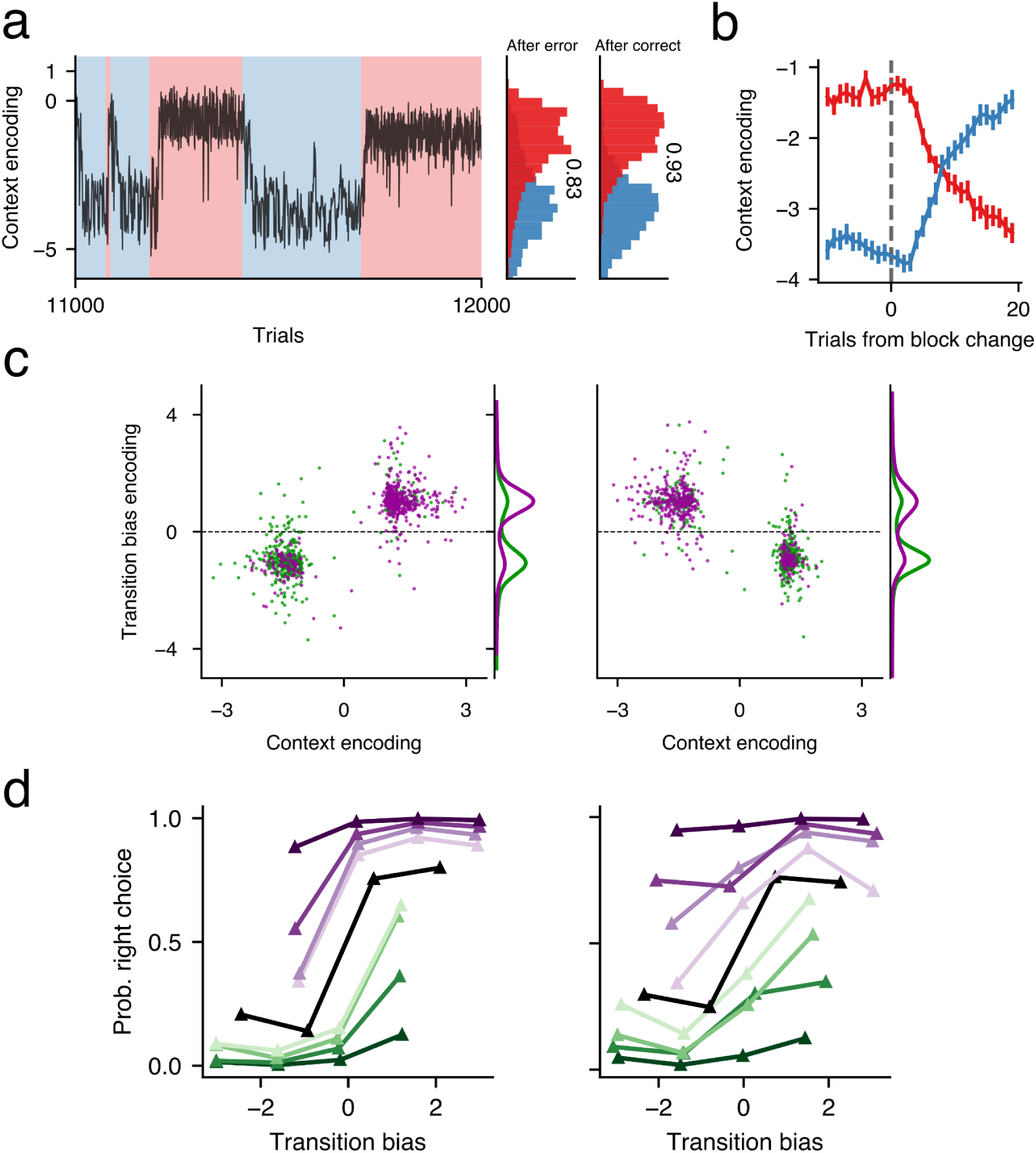
Context and transition bias encoding in 2AFC-trained networks. **a)** Dynamics of context encoding across 1000 trials spanning several blocks. **Right:** histograms for repeating (light blue) and alternating (light red) blocks show that context encoding is highly accurate. **b)** Average context encoding aligned at repeating-to-alternating (red) and alternating-to-repeating block changes (blue). **c)** Transition bias encoding versus context encoding for after-correct (left) and after-error trials (right). Only clear context trials preceded by a Left choice are shown. Dots are colored according to the true upcoming stimulus category. Each cluster shows around 20% misclassified trials corresponding to *incongruent* transitions (e.g. repetitions in an alternating block). Transition bias histograms illustrate the accuracy of the transition bias predicting the upcoming stimulus. **d)** Psychometric curves showing the network’s choice as a function of the internal encoding of the transition bias for different values of the stimulus evidence in after-correct (left) and after-error (right) trials. Color code reflects stimulus evidence (darker colors show stronger evidence; black corresponds to zero evidence). All panels correspond to the same example 2AFC-trained RNN.

**Supplementary figure 11.**
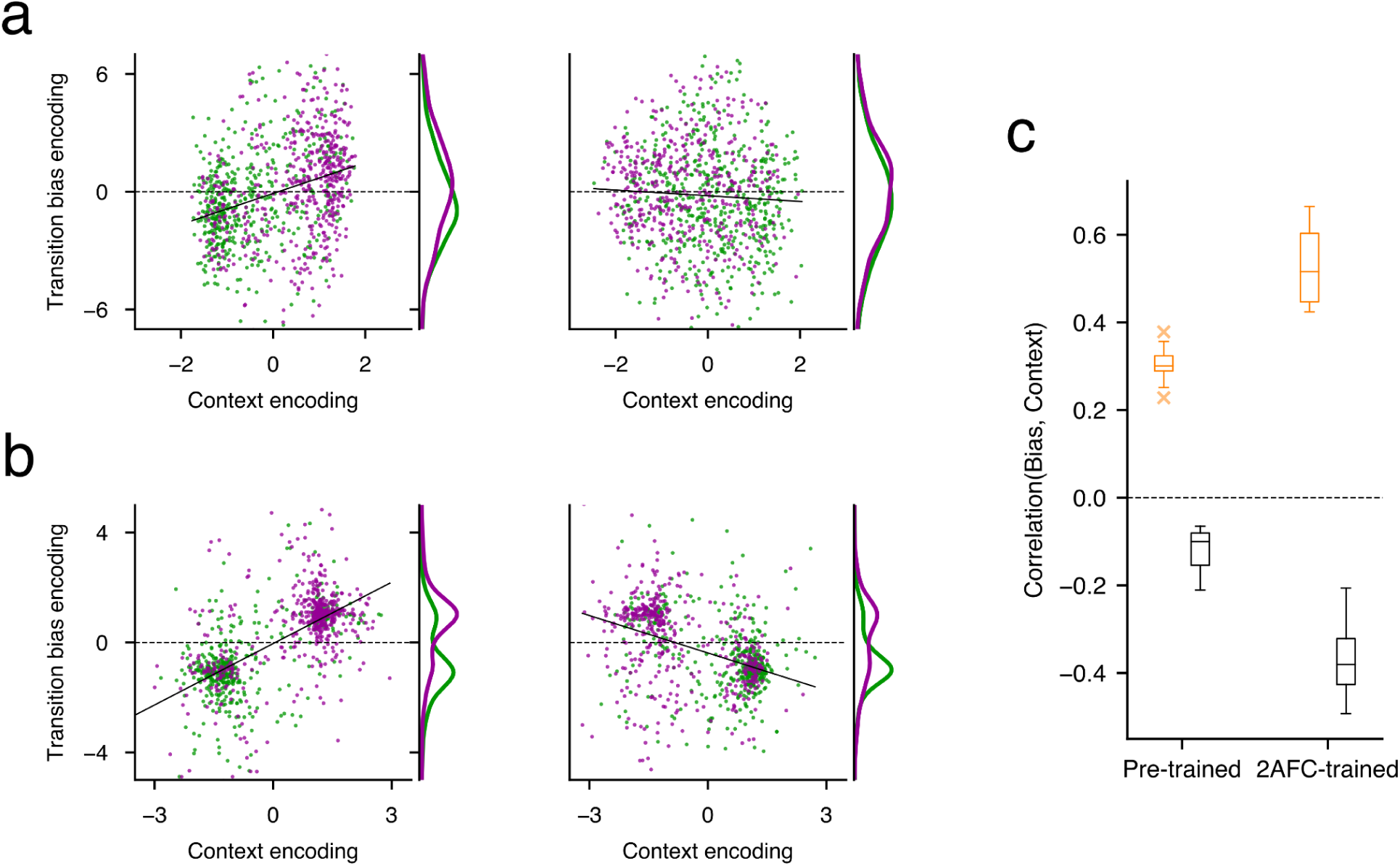
Correlation between the transition bias and the context encoding. **a, b)** Transition bias encoding versus context encoding for after-correct (left) and after-error trials (right) for the same networks shown in Fig. 6 **(a)** and Supp. Fig. 10 **(b)**. All trials preceded by a Left choice are shown. Dots are colored according to the true upcoming stimulus category. Each cluster shows around 20% misclassified trials corresponding to *incongruent* transitions (e.g. repetitions in an alternating block). Lines correspond to a linear fit. Transition bias histograms illustrate the accuracy of the transition bias predicting the upcoming stimulus. **c)** Pearson correlation coefficient between the transition bias encoding and context encoding, computed for after-correct (orange) and after-error (black) trials in pre-trained (left) and 2AFC-networks (Right).

**Supplementary figure 12.**
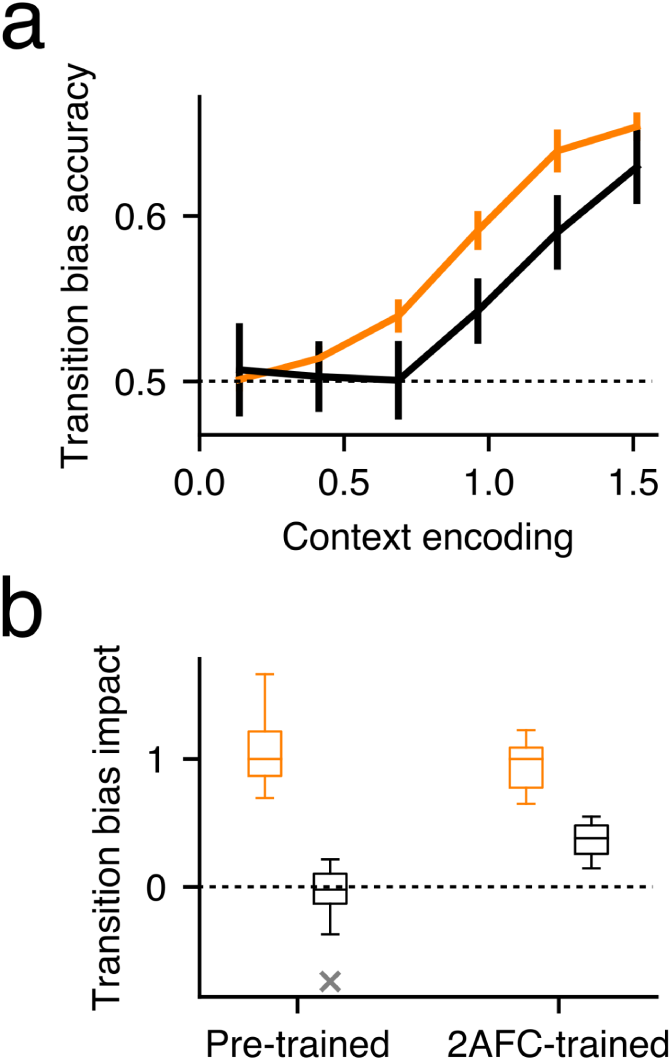
Context and transition bias encoding obtained when training the decoders with only after correct or after error trials. **a)** Transition bias accuracy versus context encoding computed separately for after-correct (orange) and after-error trials (black). Transition bias accuracy was calculated by categorizing the encoded transition bias and comparing this prediction with the true upcoming stimulus category. **b)** Normalized slopes of psychometric curves like those shown in Fig. 6h, but computed from decoders trained in only one previous outcome condition. Slopes are separately computed for pre-trained (left) and 2AFC-networks (right) for after-correct (orange) and after-error (black) trials. Slopes were normalized by the mean after-correct slope, separately for each type of RNN. Panels show the statistics from n=16 RNNs for each group. The transition bias was obtained in each condition, after-correct and after-error, from a decoder trained using only trials matching that condition (i.e. transition bias accuracy in after-correct trials was based on the decoder trained only using after-correct trials; no cross-encoding analysis was performed).

**Supplementary figure 13.**
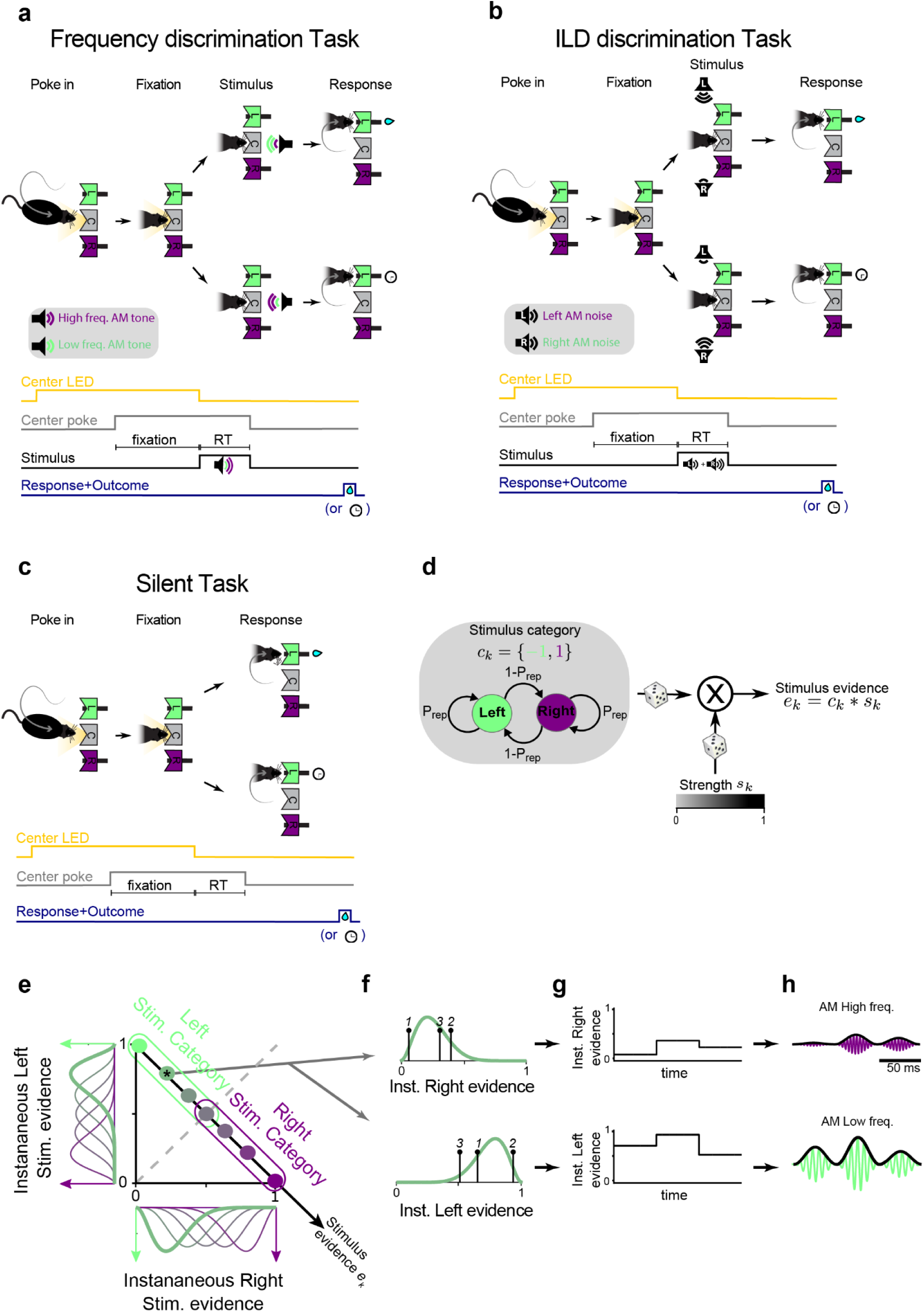
Description of the different experimental tasks used in rats. **a)** Sketch of one trial of the Frequency discrimination task: cued by the onset of the center port LED, rats poke in the center port, fixate during 300 ms and then trigger the presentation of a mixture of two AM tones, each of which is associated with reward in the left (L) or right (R) port. The dominant tone (i.e. with higher intensity) is the one that determines the reward location. Correct responses are rewarded with water, and incorrect responses are punished with a light plus a 5-s timeout. RT, reaction time. **b)** Sketch of one trial of the Interaural Level Difference discrimination task. The task structure is identical to the Frequency discrimination task, except that the stimulus consists of two AM broad-band noises, each reproduced in one of the two lateralised speakers. Rats must discriminate the loudest of the two sounds and respond seeking reward in the associated port. **c)** Sketch of one trial of the Silent task which consists of fixation (300 ms), followed by the response window triggered by the center LED offset without any sound stimulus. The trial sequence of rewarded sides for all tasks is generated using a two state Markov process (d). **d)** Schematic of the parametrization of the two-state Markov process used to generate the trial sequence of stimulus categories *c_k_* which dictates the rewarded side in the *k-*th trial (i.e. *c_k_*=1 for Righwards rewards and *c_k_*=-1 for Leftward rewards). The Markov engine is parametrized with only the probability *P_rep_* to repeat the previous category. This probability was varied in blocks of 80-100 trials between 0.8 (Repeating block) and 0.2 (Alternating block) (Fig. 1a bottom). The stimulus evidence *e_k_* is obtained in each trial from the multiplication of the stimulus category *c_k_* and the stimulus strength *s_k_*, which was independently and randomly chosen in each trial from a set of four values. (e.g. *s_k_*=0, 0.25, 0.5 and 1). **e)** Schematic of the generation of the acoustic stimuli. Each stimulus evidence *e_k_* (diagonal axis, grouped according to stim. category), is associated with two beta distributions from which the Instantaneous Left and Right stimulus evidences are drawn (see colored distributions in the left and bottom of e, respectively; color code matches the circles on the diagonal axis representing each *e_k_* ). Extreme values of the stimulus evidence *e_k_* = ±1 correspond to Dirac delta distribution centered at zero and one (see Methods for details). **f-h)** Example trial with Left stimulus category stimulus evidence *e_k_* = -0.5 showing three samples of instantaneous evidence drawn for each side (f). The samples are used to create the two temporal traces of instantaneous evidence (g). In the Frequency discrimination task (a), these traces modulate the amplitude of two AM tones with high and low frequency (h). In the ILD discrimination task (b), the instantaneous evidence traces modulate the amplitude of two AM broad-band noise bursts (not shown). The amplitude of each cycle of the AM envelope is set by the instantaneous evidence in each sample (see Methods).

**Supplementary figure 14.**
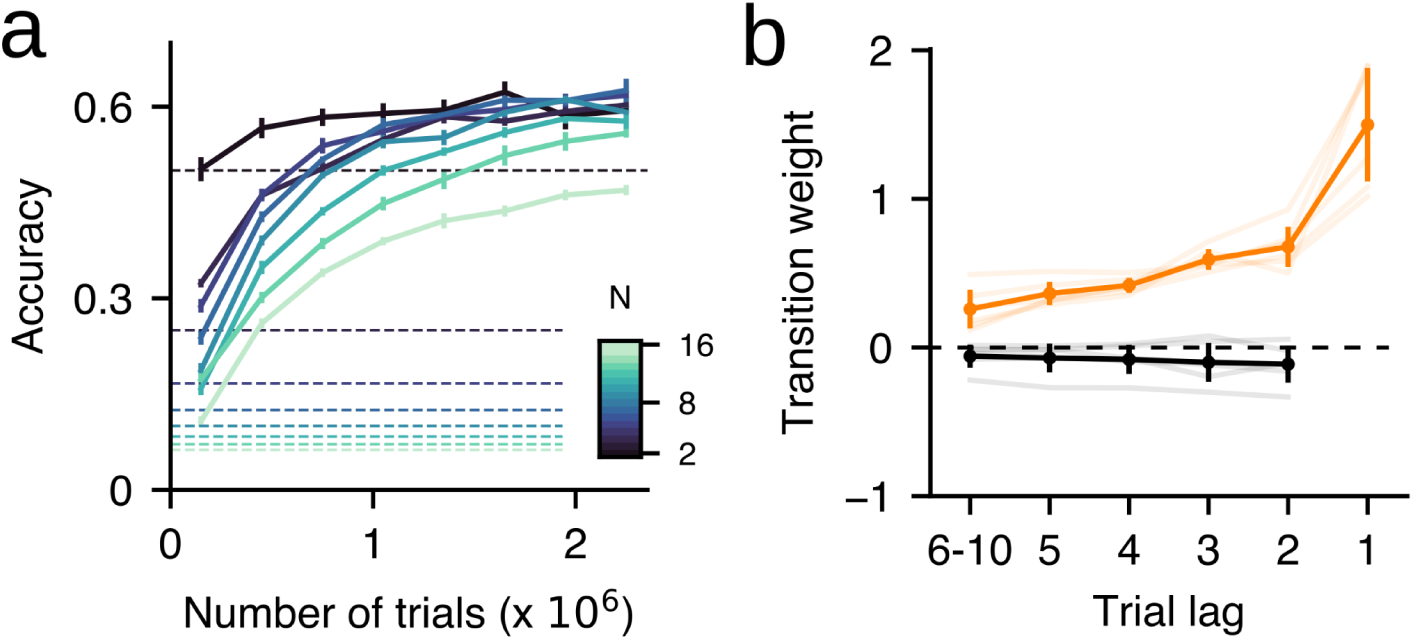
Results obtained when pre-training networks on a task with longer trials. Each trial was composed of a fixation period (3 timesteps), a stimulus lasting 4 timesteps on average (min=2, max=6) and the decision period (3 timesteps). Note that the decision period ends as soon as the network makes a decision (i.e. stops fixating and chooses one of the N possible choices). **a)** Median accuracy versus trial number in the training (n=8 RNNs) conditioned on the different number of available alternatives N=2, 4, 6, 8, 10, 12, 14, 16 (see colorbar; as in Fig. 4a). Only trials with zero stimulus evidence were used to obtain the accuracy. Dashed lines indicate the chance level (1/N) corresponding to each value of N. Error-bars correspond to standard error. **b)** GLM weights of previous correct T^++^ transitions (i.e. formed by two correct choices) computed after correct (orange) and after error (black) trials. Thick traces show average weights with s.e.m. (n=8 RNNs) whereas individual networks are shown in light traces. Error-bars correspond to standard deviation.

**Supplementary figure 15.**
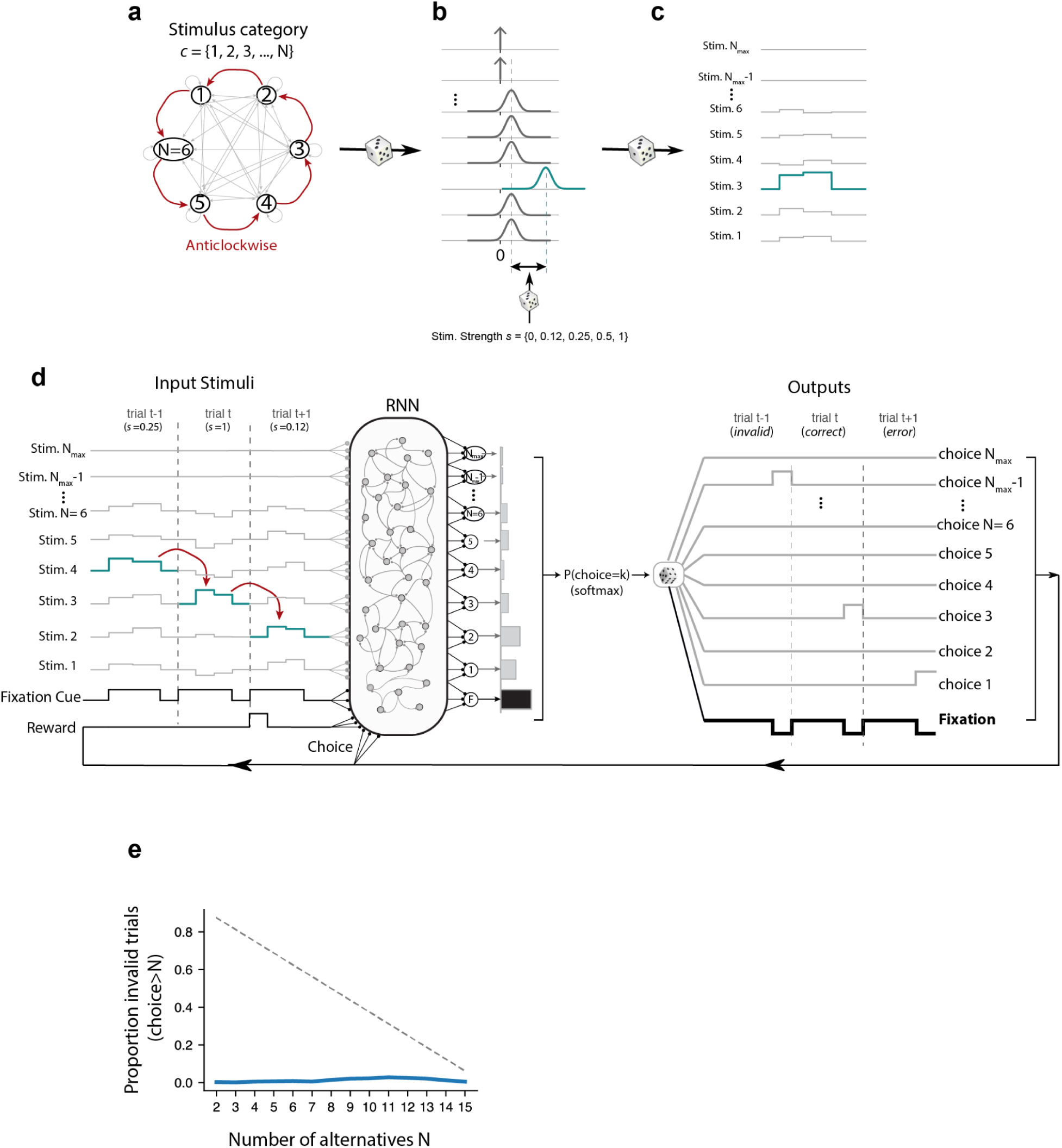
NAFC environment training. **a)** Transition diagram of the NAFC task during the ACW context when the number of valid choices was N=6. Strong red arrows represent likely transitions with probability Pr(i-1|i)=0.8 whereas gray links represent unlikely transitions with probability Pr(j|i)=0.2/(N-1)=0.04 (with j≠i+1). Using this Markov engine with N states, a sequence of stimulus categories *c* is generated. b) Normal distributions for each stimulus dimension when stimulus category the mean was μ_3_ = 0. 5 + κ *s* (blue distribution) whereas for the rest of the dimensions it was μ*_i_* = 0. 5 − κ *s* , with  *i* ∈ [1, 2, 4, … *N*] (gray Gaussian distributions). The stimulus strength *s* is randomly drawn in each trials from the set of discrete values . The prefactor κ only depends on N ∈ [0 , 0. 12, 0. 25, 0. 5, 1]. The prefactor κ only depends on N (see Stimulus section in Methods). The distribution of the dimensions corresponding to invalid choices above N (i.e. 7, 8, … N_max_) was a Dirac delta centered at zero (i.e. all drawn values were zero). **c)** Example stimuli drawn from the distributions shown in (b) composed of two independent and consecutive samples for each dimension. **d)** At each timestep, the networks received as input a fixation cue, the N_max_ input stimuli, and the reward and action at the previous timestep. The output was obtained from (N_max_+1) linear readout units that represented each of the possible outputs. At each timestep, the activity of the readout units was passed through a multivariate softmax function in order to choose between fixation, or one of the N_max_ possible choices. The number of valid choices, N, was changed every 5k trials between 2 and N_max_. The example shows a sequence of 3 trials following a pattern consistent with the ACW context. In response, the network selects an invalid choice (N_max_-1) in trial *t*-1, the correct choice (3) in trial *t* and an incorrect choice (1) in trial *t*+1 . **e)** Average proportion of invalid choices (i.e. above N) during training as a function of N for networks trained with N_max_ = 16 (blue trace). The dashed line indicates the proportion that would be expected for an agent making random choices.

